# Pallidal prototypic neuron and astrocyte activities regulate flexible reward-seeking behaviors

**DOI:** 10.1101/2025.02.10.637554

**Authors:** Shinwoo Kang, Minsu Abel Yang, Aubrey Bennett, Seungwoo Kang, Sang Wan Lee, Doo-Sup Choi

**Author notes:** Corresponding authors (Doo-Sup Choi). These authors contributed equally to this work as the first author.

## Abstract

Behavioral flexibility allows animals to adjust actions to changing environments. While the basal ganglia are critical for adaptation, the specific role of the external globus pallidus (GPe) is unclear. This study examined the contributions of two major GPe cell types—prototypic neurons projecting to the subthalamic nucleus (Proto^GPe→STN^ neurons) and astrocytes—to behavioral flexibility. Using longitudinal operant conditioning with context reversals, we found that Proto^GPe→STN^ neurons dynamically represent contextual information correlating with behavioral optimality. In contrast, GPe astrocytes exhibited gradual contextual encoding independent of performance. Deleting Proto^GPe→STN^ neurons impaired adaptive responses to changing action-outcome contingencies without altering initial reward-seeking acquisition, highlighting their specific role in enabling behavioral flexibility. Furthermore, we discovered that Proto^GPe→STN^ neurons integrate inhibitory striatal and excitatory subthalamic inputs, modulating downstream basal ganglia circuits to support flexible behavior. This research elucidates the complementary roles of Proto^GPe→STN^ neurons and astrocytes in cellular mechanisms of flexible reward-seeking behavior.

## Introduction

Animals continuously adapt their strategies to align with environmental changes^1, 2^, which necessitates behavioral flexibility. This adaptability is essential for survival, allowing animals to modify their actions when previously learned behaviors no longer achieve the desired outcomes. Longitudinal experimental designs incorporating context reversals^3–7^ are particularly suited to studying the mechanisms underlying behavioral flexibility, as they capture the acquisition, stabilization, and subsequent adaptations of behavioral strategies in response to altered contingencies. These approaches also enable examining the dynamic cellular activity associated with flexible behavior^3–5^.

The basal ganglia are central to behavioral flexibility^6, 8–10^, integrating information related to context^11^, reward^12–14^, and motor planning^15, 16^ to support adaptive responses in changing environments. While the involvement of basal ganglia in behavioral flexibility is well-established, much of the research in this area has predominantly focused on the role of the striatum, particularly its different subregions^17–19^ and cell types^20, 21^. Studies have extensively investigated striatal circuits and plasticity mechanisms^21, 22^ underlying various aspects of flexible behavior, including action selection, reversal learning, and cognitive control^23, 24^. However, the specific contributions of other key basal ganglia nuclei to behavioral flexibility remain comparatively less explored.

Specifically, the external globus pallidus (GPe) functions as a critical hub within the basal ganglia, processing inputs from the cortex, thalamus, brainstem, and dorsal striatum and projecting to key structures such as the subthalamic nucleus (STN), substantia nigra pars reticulata (SNr), and the parafascicular (Pf) nucleus of the thalamus. Through reciprocal feedback loops with the dorsal striatum, the GPe regulates basal ganglia activity, facilitating both motor and cognitive processes^25, 26^.

Traditionally, the GPe, the STN, and the SNr have been viewed within a ’canonical information flow’ model of the basal ganglia indirect pathway (Striatum→GPe→STN→SNr), where GPe output inhibits the STN, and the STN, in turn, excites the SNr^27, 28^. Recent studies have expanded this view of the GPe from a passive relay^27, 29, 30^ to an active participant in sensory integration, reward processing, and learning^10, 31–33^. Importantly, substantial recent evidence challenges this simplified ’canonical information flow’ picture, suggesting a more complex, ’non-canonical’ organization. This includes evidence for direct GPe→SNr projections^10, 34^, cortico-STN ’hyperdirect’ pathway^35–38^, and feedback circuitry from the STN to the GPe (STN→GPe)^10, 39, 40^, implying that the GPe may receive excitatory input from the STN and directly modulate SNr output, challenging the traditional GPe-STN inhibitory relay.

Within the GPe, neurons and astrocytes cooperate for such functions. Particularly, prototypic neurons, which comprise over half of GPe neurons^34, 41^, integrate inhibitory inputs from indirect medium spiny neurons (iMSNs) in the dorsal striatum and excitatory signals from the STN, while also forming local inhibitory connections^39, 40^. These neurons are connected to multiple basal ganglia and thalamic targets^10, 34, 42^, shaping basal ganglia output to influence motor and cognitive functions^10, 32, 39, 43, 44^. On the other hand, astrocytes, the most abundant cell type in the GPe^45^, play complementary roles by maintaining synaptic function^46–50^ and modulating neural circuits^46, 49, 51–54^. Together, prototypic neurons and astrocytes shape GPe activity, contributing to the broader dynamics of the basal ganglia^26, 34^.

Here, we investigated the contributions of two major GPe cell types, prototypic neurons projecting to the STN (Proto^GPe→STN^ neurons) and astrocytes (GFAP+ cells), to behavioral flexibility during repetitive reward-seeking conditioning. Through a longitudinal operant conditioning task with context reversals, we first demonstrated that Proto^GPe→STN^ neurons, not astrocytes, develop contextual encoding as learning progresses, and this encoding capability correlates with behavioral optimality. Notably, GPe astrocytes displayed gradual contextual encoding that was not correlated with behavioral performance. Caspase-dependent ablation of Proto^GPe→STN^ neurons revealed that initial reward-seeking acquisition was unaffected, but adaptive responses to changing action-outcome contingencies were significantly impaired, directly demonstrating their specific role in enabling behavioral flexibility. Optogenetic manipulation of iMSNs in the dorsolateral striatum (DLS) revealed that striatal inputs provided critical information for Proto^GPe→STN^ neurons during adaptation. To test the contrasting predictions of the canonical versus non-canonical information flow models, we examined the effect of chemogenetic manipulation of STN neurons on behavioral flexibility. Chemogenetic experiments showed that STN neurons relayed context-change information to Proto^GPe→STN^ neurons, which modulated downstream motor circuits *via* direct projections to the SNr to support flexible behavior through axonal collateral^10, 34^. Our findings demonstrate the necessity of GPe for behavioral flexibility, where Proto^GPe→STN^ neurons integrate inputs from the striatal and subthalamic nucleus, contributing to the modulation of basal ganglia circuitry supports adaptive behavior.

## Results

### Longitudinal calcium imaging of reward- and reversal-seeking behaviors

Given the emerging evidence of the GPe’s role in modulating motor and non-motor functions^10, 31–33, 39, 43, 44, 55^, we sought to investigate its differential contributions to the acquisition of reward-seeking behavior and adaptation to changing reward contingencies. To address this, we employed a fixed ratio 1 (FR1) operant conditioning with a reversal-learning task. Mice were trained on an FR1 schedule, where a single nose-poke in the active hole (active nose-poke, ANP) delivered a reward (10 μl of 20% sucrose) accompanied by light and sound cues in an operant conditioning chamber for 54 days (Fig. 1a). Rewards were retrieved from the magazine following each successful ANP (Fig. 1b). Each daily session lasted either 60 minutes or until mice completed 60 active nose-pokes.

**Figure 1.**
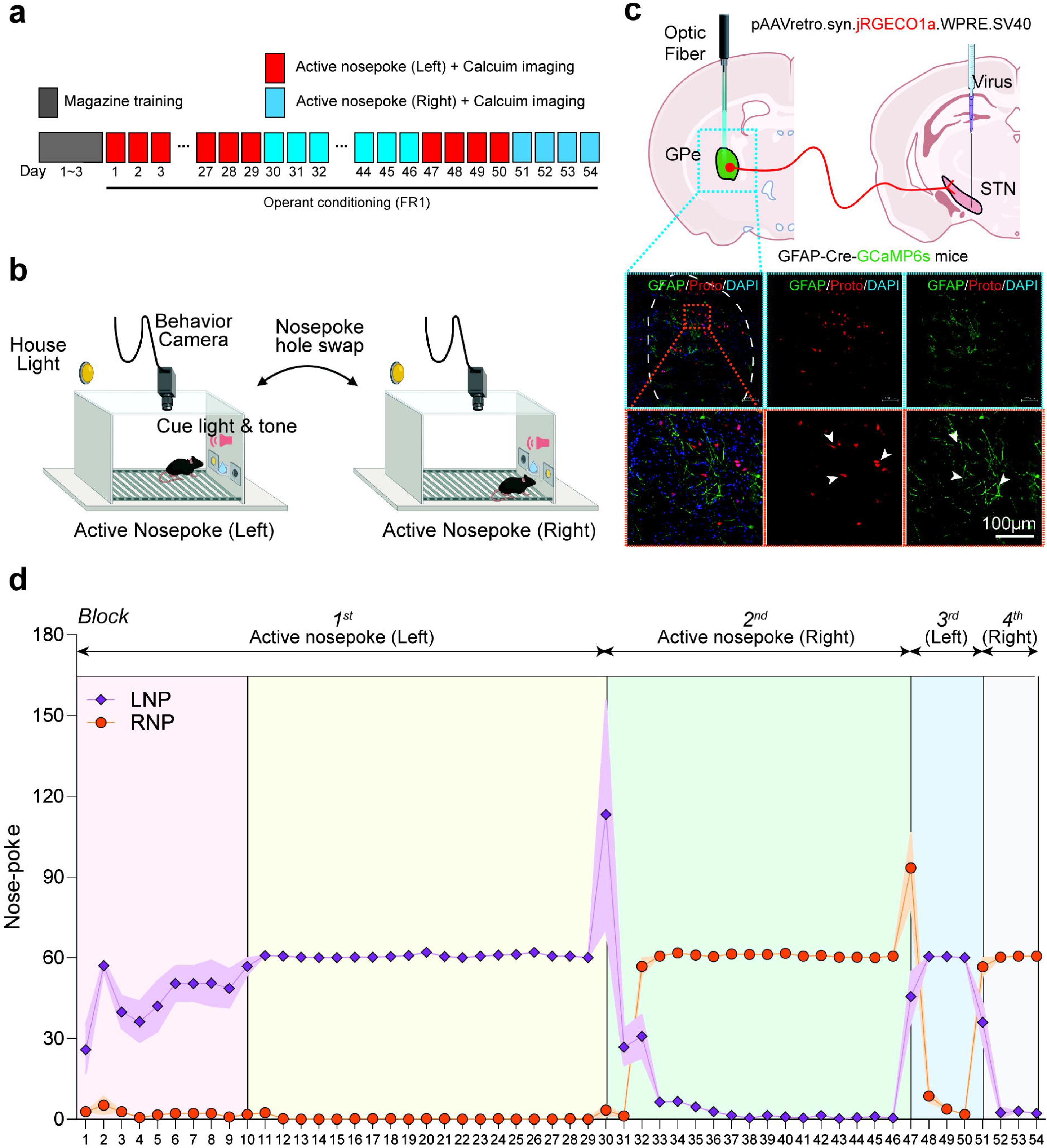
Operant conditioning task and fiber photometry. **a,** Schematic of the FR1 task training schedule. **b,** Schematic of operant conditioning task with reversal. During the left block, the active (rewarded) nose-poke is the left nose-poke. On the other hand, during the right block, the active nose-poke is the right nose-poke. **c,** Fiber photometry configuration to measure *in vivo* Ca^2+^ dynamics and histological validation of GPe astrocytes (green, GCaMP6s) and Proto^GPe→STN^ neurons (red, jRGECO1a). Scale: 100 µm. **d,** Nose-poking behaviors during the FR1 task training. Over the course of the 54-day experiment, the context was reversed three times (left, right, left, right): Day 30, Day 47, and Day 51. This design resulted in four distinct blocks: Days 1-29 (1^st^ block), Days 30-46 (2^nd^ block), Days 47-50 (3^rd^ block), and Days 51-54 (4^th^ block). Shaded areas represent SEM. Data consist of a total of 261 sessions, collected across four blocks from 5 mice: *n =* 140 (1^st^ block), 82 (2^nd^ block), 19 (3^rd^ block), and 20 (4^th^ block). See Supplementary Table 1 for full statistical information.

To evaluate behavioral flexibility, we implemented a reversal-learning task (Fig. 1b) using two distinct contexts: a left context (Fig. 1b, left) and a right context (Fig. 1b, right). In the left context, the left nose-poke was the ANP delivering a reward, while the right nose-poke was non-rewarded (inactive, INP). Conversely, the roles were reversed in the right context, with the right nose-poke designated as the ANP and the left nose-poke as the INP. For each block, the context remained stable, and the location of the active hole did not change. Without explicit cues, the context switched abruptly, reversing the location of the active and inactive nose-poke holes. Throughout the 54-day experiment, the context was reversed three times: starting with the left context on Day 1, switching to the right on Day 30, reverting to the left context on Day 47, and finally returning to the right context on Day 51. This design resulted in four distinct blocks: Days 1-29 (Fig. 1d, 1^st^ block, left context), Days 30-46 (Fig. 1d, 2^nd^ block, right context), Days 47-50 (Fig. 1d, 3^rd^ block, left context), and Days 51-54 (Fig. 1d, 4^th^ block, right context). This protocol allowed us to examine how mice adapt to sudden changes in reward contingencies across multiple reversals.

To simultaneously monitor GPe astrocytes and prototypic neurons during behavior, we employed fiber photometry in GFAP-Cre/DIO-GCaMP6s mice. Proto^GPe→STN^ neurons were selectively labeled by injecting a retrograde adeno-associated virus (AAV) driven by a synapsin promoter, expressing the fluorescent calcium indicator jRGECO1a, into the STN of GFAP-Cre/DIO-GCaMP6s mice (Fig. 1c). This dual-labeling strategy, combined with time-division multiplexing and isosbestic control, enabled us to capture activity from both astrocytes and Proto^GPe→STN^ neurons simultaneously in the GPe, with minimal signal crosstalk. While mice performed the reversal-learning task over 54 days (*n* = 261 sessions from 5 mice, Fig. 1d and Extended Data Fig. 1), we recorded astrocytic and neuronal activity daily, providing longitudinal insights into their dynamic roles in behavioral adaptation.

### Dynamic interaction of GPe astrocytes and Proto^GPe^**^→^**^STN^ neurons during learning and behavioral adaptation

Using longitudinal FR1 experiments, we examined how distinct cellular populations in the GPe — astrocytes and Proto^GPe→STN^ neurons — respond during different phases of task learning, including the stabilization of learned behaviors and adaptation to reversals in action strategies. To confirm that mice acquired the necessary task policies across different blocks, we utilized DeepLabCut^56^ to track their position and pose from video recordings (Fig. 2a). Position data were analyzed alongside the timing of key action events, including left nose-poke (LNP), right nose-poke (RNP), magazine entry (ME), and magazine exit (MX). This data was used to construct transition probability maps between events, providing a comprehensive view of behavioral patterns across learning phases (Fig. 2b).

**Figure 2.**
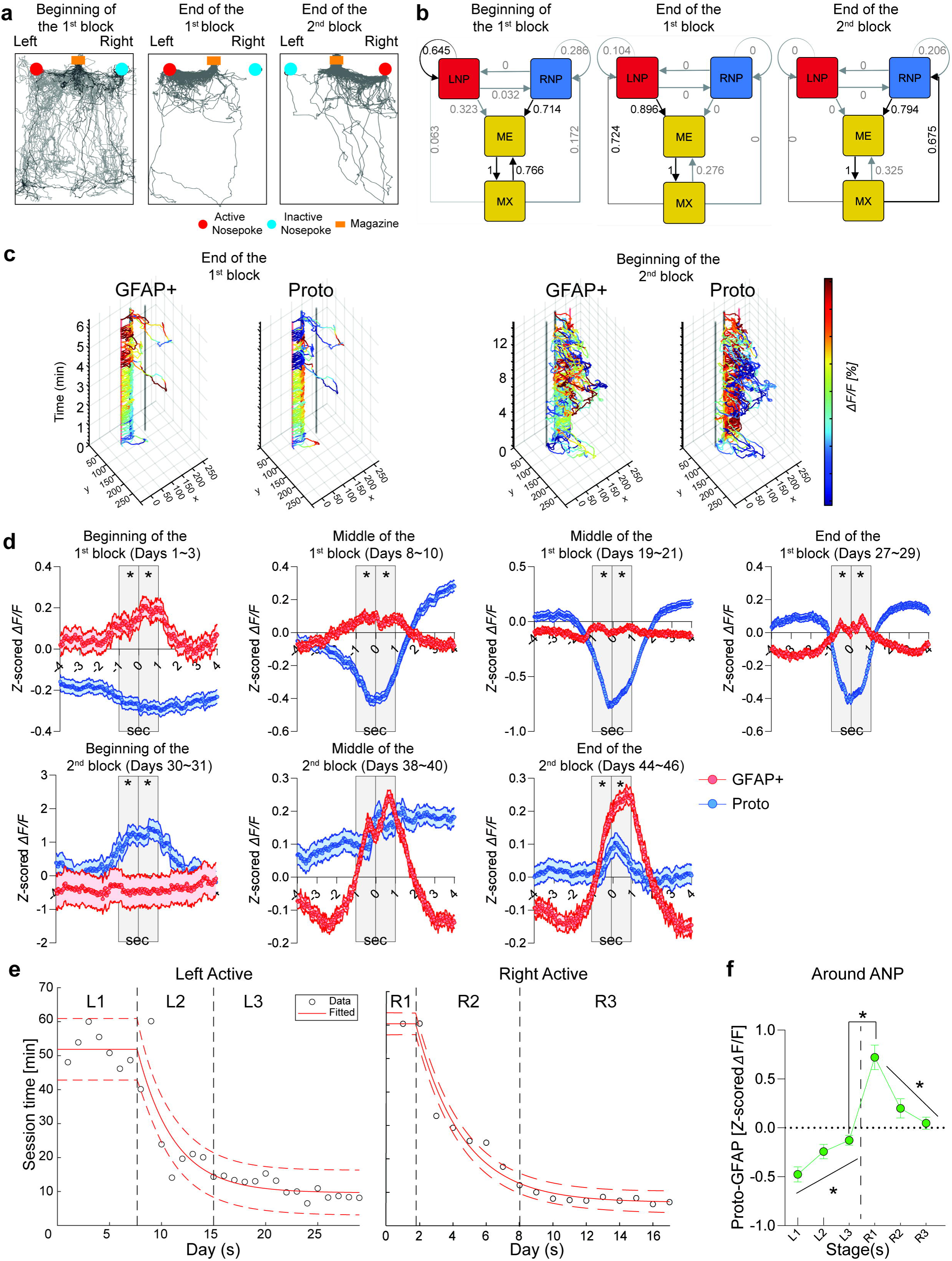
Dynamics of GPe astrocytes and Proto^GPe→STN^ neurons during a longitudinal operant conditioning task with context reversal. **a-b,** Example behavioral data at different learning stages. Beginning of the 1^st^ block (Day 1), End of the 1^st^ block (Day 29), and End of the 2^nd^ block (Day 46) from the left. **a,** Trajectory plots showing the example mouse movement in the operant conditioning chamber during an entire session. Key apparatus components related to different action events are depicted: active nose-poke hole (red), inactive nose-poke hole (blue), and the magazine (yellow). **b,** Transition probability maps between action events. For each state, the black line indicates transition with the highest probability, while gray lines represent lower probability transitions. **c,** Representative calcium traces recorded from GPe astrocytes (GFAP+) and Proto^GPe→STN^ neurons (Proto) right before (Day 29) and after (Day 30) the first context reversal. **d,** Averaged calcium signals of GPe astrocytes and Proto^GPe→STN^ neurons aligned to the active nose-poke at 0 s. Shaded areas represent SEM. * denotes when the average GPe astrocyte activity during the specified interval is significantly different from the average Proto^GPe→STN^ neuron activity (*P* < 0.05, Paired t-test). **e,** Session-time-based learning phase division. Left for the 1^st^ block (Left active nose-poke). Right for the 2^nd^ block (Right active nose-poke). **f,** Longitudinal calcium imaging revealed increasing differences in Ca^2+^ activity between GPe astrocytes and Proto^GPe→STN^ neurons during learning (L1-L3), with a peak observed following the first reversal (R1). This activity difference gradually decreased as animals adapted to the new context (R1-R3). Data are represented as mean ± SEM, except for **f**, where data are represented as estimated marginal mean and standard error from the linear mixed-effects model; Data consist of a total of 219 sessions, collected across two blocks from 5 mice: *n* = 32 (L1), 34 (L2), 74 (L3), 12 (R1), 19 (R2), and 48 (R3). See Supplementary Table 1 for full statistical information.

At the beginning of the first block, mice exhibited exploratory behavior, interacting indiscriminately with both nose-poke holes and frequently moving along the chamber edges, areas irrelevant to task performance (Fig. 2a, beginning of the 1^st^ block). During this phase, mice often transitioned from MX to ME without preceding ANP, resulting in no reward (Fig. 2b, beginning of the 1^st^ block). By the end of the first block, mice shifted to exploitative behavior, selectively approaching the active left nose-poke hole and alternating consistently between the left nose-poke hole and the magazine for reward acquisition (Fig. 2a, end of the 1^st^ block). This behavior formed a stable sequence: LNP→ME→MX→LNP (Fig. 2b, end of the 1^st^ block). Following the reversal, at the end of the second block, mice adapted by switching to RNP, which is designated as active, and established a similar exploitative pattern. This adaptive behavior was reflected in the transition loop between RNP, ME, and MX (Fig. 2a-b, end of the 2^nd^ block). These findings highlight the dynamic changes in behavior as mice transitioned from exploration to exploitation and adapted to reversed contingencies.

After confirming successful learning across blocks, we analyzed the activity of GPe astrocytes and Proto^GPe→STN^ neurons immediately following the reversal. Prior to the reversal, astrocytes exhibited elevated Ca^2+^ activity around ANP compared to Proto^GPe→STN^ neurons (Fig. 2c left, end of the 1^st^ block). However, immediately after the reversal, Proto^GPe→STN^ neurons showed a marked increase in Ca^2+^ activity, surpassing that of astrocytes (Fig. 2c right, beginning of the 2^nd^ block).

To further investigate these dynamics, we analyzed calcium signals within an 8-second window, spanning 4 seconds before and 4 seconds after ANP (Fig. 2d and Extended Data Fig. 2). During the early-stage of initial learning (Days 1–3), astrocytes exhibited higher activity around ANP compared to Proto^GPe→STN^ neurons (Fig. 2d, beginning of the 1^st^ block and see also the video recording integrated with cell-specific activity plot, Extended video 1). This trend persisted as learning progressed within the first block (Fig. 2d, 1^st^ row). In contrast, immediately following the reversal, Proto^GPe→STN^ neuron activity increased significantly, surpassing that of astrocytes (Fig. 2d, beginning of the 2^nd^ block, see the video recording in Extended video 2). As learning progressed and mice adapted to the new context, the disparity between Proto^GPe→STN^ neuron activity and astrocyte activity gradually diminished. By the middle of the second block, the activity levels of Proto^GPe→STN^ neurons and astrocytes were approximately equal (Fig. 2d, middle of the 2^nd^ block). Toward the end of the second block, astrocytes again displayed greater activity than Proto^GPe→STN^ neurons, re-establishing the initial trend observed during the first block (Fig. 2d, end of the 2^nd^ block, see the video recording in Extended video 3).

To analyze the evolution of Ca^2+^ activity differences between astrocytes and Proto^GPe→STN^ neurons during learning — while accounting for differences in learning speeds — we divided each block into three distinct phases for each subject (Fig. 2e): (1) pre-learning (L1 in the 1^st^ block, R1 in the 2^nd^ block), (2) learning (L2 in the 1^st^ block, R2 in the 2^nd^ block), and (3) post-learning (L3 in the 1^st^ block, R3 in the 2^nd^ block). For each phase, we calculated the average difference in Ca^2+^ activity between Proto^GPe→STN^ neurons and astrocytes within a 4-second window, spanning 2 seconds before and after ANP. The results showed a significant increase in the average Ca^2+^ activity difference as learning progressed within the first block, from pre-learning to post-learning phases (Fig. 2f, L1-L3). Immediately after the reversal, this difference exhibited a sharp increase (Fig. 2f, L3-R1). As learning advanced in the second block, transitioning from pre-learning to post-learning, the average difference gradually diminished (Fig. 2f, R1-R3).

These findings reveal that astrocytes and Proto^GPe→STN^ neurons show varying Ca^2+^ activity levels during the learning and adaptation phases, which change over time alongside shifts in task performance. Moreover, the observed activity patterns differ between the first and second blocks, indicating that both astrocytes and Proto^GPe→STN^ neurons exhibit Ca^2+^ signals, which depend on the subject’s learning phase and context.

### GPe astrocytes and Proto^GPe^**^→^**^STN^ neurons develop contextual encoding differently as learning progresses

For flexible behavior, animals need to adjust their action based on the broader situational context rather than solely on isolated events. This process is supported by contextual encoding, which enables the brain to generalize from past experiences and adapt behavior to new contexts^57, 58^. We hypothesized that cellular populations in the GPe, specifically Proto^GPe→STN^ neurons and astrocytes, contribute to facilitating flexible behavior through contextual encoding.

To test this hypothesis, we investigated whether GPe astrocytes (Fig. 3a top) and Proto^GPe→STN^ neurons (Fig. 3a bottom) differently represent action events depending on the *status quo* context. We specifically analyzed Ca^2+^ activity surrounding ME and MX events, as these actions consistently involve interaction with the magazine—the site of reward delivery—regardless of the context. Unlike left or right nose-poke actions, which yield context-dependent outcomes based on the active contingency, ME and MX events provide context-invariant interactions. This distinction allowed us to specify the capability of GPe cells to contextual encoding without the confounding effects of changing reward contingencies, offering a stable reference point for analyzing context representation.

**Figure 3.**
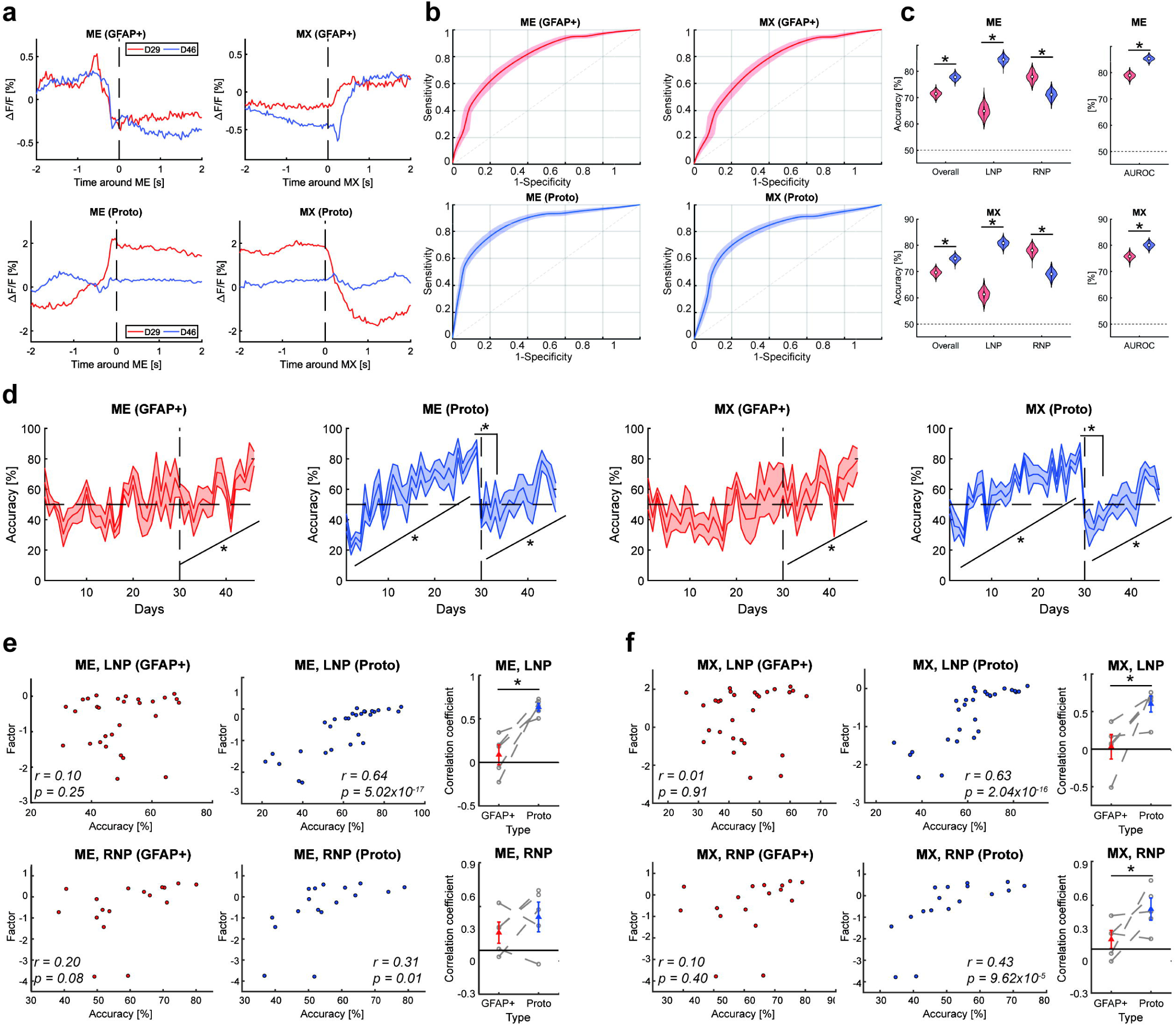
Contextual encoding of GPe astrocytes and Proto^GPe→STN^ neurons and their associations with the behavioral optimization. **a,** Example Ca^2+^ trace of GPe astrocytes (GFAP+, top) and Proto^GPe→STN^ neurons (Proto, bottom) aligned to the magazine-related action events. Left for Ca^2+^ activity around ME. Right for Ca^2+^ activity around MX. Blue line for average activity at Day 29 (D29), the end of the 1^st^ block (left context). Red line for average activity at Day 46 (D46), the end of the 2^nd^ block (right context). **b-c,** Classifier performance. Two classifiers were trained on Ca^2+^ activity data from 5 mice: the ME classifier was trained with *n =* 1307 (1^st^ block) and 975 (2^nd^ block) traces, and the MX classifier was trained with *n* = 1316 (1^st^ block) and 985 (2^nd^ block) traces. **b,** ROC curves depicting the predictive performance of Ca^2+^ trace around behavioral events for contextual encoding. Top and bottom for GPe astrocytes (GFAP+) and Proto^GPe→STN^ neurons (Proto), respectively. Left and right for ME and MX, respectively. **c,** Comparison of classifier performance measures across cell types. Metrics include accuracy across blocks (Overall), accuracy in the left context (LNP), accuracy in the right context (RNP), and AUROC from the left. Top and bottom for ME and MX, respectively. **d,** Change in classification accuracy across days. Classifier trained on Ca^2+^ trace of GPe astrocytes around ME [ME (GFAP+)], classifier trained on Ca^2+^ trace of Proto^GPe→STN^ neurons around ME [ME (Proto)], classifier trained on Ca^2+^ trace of GPe astrocytes around MX [MX (GFAP+)], and classifier trained on Ca^2+^ trace of Proto^GPe→STN^ neurons around MX [MX (Proto)] from the left. Classification accuracy analysis is based on data from 5 mice, across a total of 221 sessions: *n =* 139 (1^st^ block), and 82 (2^nd^ block). **e-f,** Correlation between degree of contextual encoding (accuracy) and behavioral optimality (factor). GPe astrocytes (GFAP+), Proto^GPe→STN^ neurons (Proto), and comparison of correlation coefficients between two cell types from the left. Top and bottom for the first (left context, LNP) and second block (right context, RNP), respectively. Correlation analysis is based on data from 5 mice, across a total of 220 sessions: *n =* 138 (1^st^ block), and 82 (2^nd^ block). **e,** Activity around ME. **f,** Activity around MX. Data are represented as mean ± SEM, except for violin plots in **c**. See Supplementary Tables 1-3 for full statistical information.

To investigate this, we performed decoding analyses on Ca^2+^ activity recorded during the final five sessions of the first block (Fig. 3a red lines, left context, Days 25-29) and second block (Fig. 3a blue lines, right context, Days 42-46), when learning had stabilized. A linear support vector machine (SVM) classifier was trained to predict the contextual state (left or right) based on Ca^2+^ activity surrounding ME and MX events. Classifiers trained on astrocyte Ca^2+^ activity surrounding both ME (Fig. 3b top left) and MX events (Fig. 3b top right) performed significantly above chance, as measured by the area under the receiver operating characteristic curve (AUROC, ME: 78.88%; MX: 75.71%) and three accuracy metrics (ME: Fig. 3c top, red violins, MX: Fig. 3c bottom, red violins). The context-specific accuracy metrics (Fig. 3c) include: (1) accuracy across blocks (Overall), indicating the proportion of Ca^2+^ activities correctly classified according to their respective context; (2) accuracy on the left context (LNP), showing the proportion of Ca^2+^ activities from the first block (left context) that were correctly classified; and (3) accuracy on the right context (RNP), showing the proportion of Ca^2+^ activities from the second block (right context) that were correctly classified. Similarly, classifiers based on Proto^GPe→STN^ neuronal activity surrounding ME (Fig. 3b bottom left) and MX events (Fig. 3b bottom right) also performed significantly above chance for AUROC (ME: 85.32%; MX: 80.04%) and all three-accuracy metrics (ME: Fig. 3c top, blue violins, MX: Fig. 3c bottom, blue violins). Importantly, to ensure the robustness of these classification results, we validated the SVM performance across different input time windows, confirming that significant classification accuracy was maintained for a range of window durations (Supplementary Table 2, see Methods for details).

Interestingly, Proto^GPe→STN^ neurons demonstrated higher accuracy across blocks and AUROC compared to astrocytes for both ME (Fig. 3c top) and MX events (Fig. 3c bottom), indicating stronger contextual encoding. Specifically, Proto^GPe→STN^ neurons exhibited superior accuracy in the first block for both ME (Fig. 3c top, LNP) and MX events (Fig. 3c bottom, LNP), effectively classifying Ca^2+^ activities associated with the left context. Conversely, astrocytes outperformed Proto^GPe→STN^ neurons in the second block for both ME (Fig. 3c top, RNP) and MX events (Fig. 3c bottom, RNP), showing greater classification performance for Ca^2+^ activities recorded during the right context.

To quantify the degree of contextual encoding in individual Ca^2+^ traces, we employed trained classifiers to discriminate contexts based on GPe astrocytes and Proto^GPe→STN^ neuronal activity (Fig. 3b-c). Classification accuracy, as detailed in the ‘Classification analysis’ of the Methods, served as a quantitative measure of this contextual encoding. For each session, we measured the classification accuracy - the probability of correctly identifying the session’s context from Ca^2+^ activity – and analyzed its change across days within the first and second blocks.

Our analysis revealed distinct patterns of contextual encoding between GPe astrocytes and Proto^GPe→STN^ neurons, highlighting their unique roles in response to learning and context changes (Fig. 3d and Extended Data Fig. 3). GPe astrocytes exhibited rather constant classification accuracy for both ME (Fig. 3d 1^st^ column, Days 1-29) and MX events (Fig. 3d 3^rd^ column, Days 1-29) throughout the first block, indicating a consistent encoding pattern that was unaffected by ongoing learning. Furthermore, comparisons of classification accuracy immediately before and after the reversal showed no significant changes, suggesting consistent encoding across context shifts (ME: Fig. 3d 1^st^ column, MX: Fig. 3d 3^rd^ column, Days 29-30). However, during the second block, classification accuracy for astrocytes increased significantly as learning progressed (ME: Fig. 3d 1^st^ column, MX: Fig. 3d 3^rd^ column, Days 30-46). This suggests that astrocytes reinforce contextual encoding following the reversal, supporting stability as new learning becomes consolidated.

In contrast, Proto^GPe→STN^ neurons exhibited a more dynamic pattern of contextual encoding, with classification accuracy reflecting temporal changes in their encoding capabilities. During the first block, classification accuracy for both ME (Fig. 3d 2^nd^ column, Days 1-29) and MX events (Fig. 3d 4^th^ column, Days 1-29) increased significantly, indicating a progressive refinement of contextual encoding as learning advanced. Notably, classification accuracy dropped sharply following the context reversal, highlighting a temporary disruption in encoding when outcome contingencies shifted (ME: Fig. 3d 2^nd^ column, MX: Fig. 3d 4^th^ column, Days 29-30). This suggests that the previous contextual encoding of Proto^GPe→STN^ neurons no longer aligned with the new context, necessitating re-encoding. As learning progressed during the second block, classification accuracy for Proto^GPe→STN^ neurons increased again, reflecting a recalibration to the new context and demonstrating their capability for flexible adaptation (ME: Fig. 3d 2^nd^ column, MX: Fig. 3d 4^th^ column, Days 30-46).

These findings suggest that Proto^GPe→STN^ neurons play a dynamic role, adjusting their encoding to accommodate shifting task demands. This dynamic adjustment likely contributes to behavioral flexibility. In contrast, GPe astrocytes primarily represent contexts that are more stable compared to Proto^GPe→STN^ neurons, particularly during initial learning.

### Contextual encoding of Proto^GPe^**^→^**^STN^ neurons correlates with behavioral optimality

Building on the observation that both Proto^GPe→STN^ neurons and GPe astrocytes exhibit contextual encoding, we sought to further clarify the specific roles of these two cell types in behavioral flexibility. Specifically, we investigated whether the degree of contextual encoding in each cell type correlates with the behavioral optimality of the corresponding mouse. Behavioral optimality, which reflects the ability to achieve efficient and accurate performance in a changing environment, serves as a proxy for behavioral flexibility. We hypothesized that a positive correlation between the degree of contextual encoding in one cell type and behavioral optimality would suggest a strong association between that cell type and behavioral flexibility.

To quantify behavioral optimality, we employed three behavioral metrics that capture different aspects of optimal performance in the FR1 task. First, we measured session time, as optimal behavior is expected to result in quicker session completion (Extended Data Fig. 1c). Second, we calculated the discrimination ratio, which quantifies the extent to which animals preferentially select the active nose-poke hole over the inactive one (Extended Data Fig. 1a). Third, we analyzed trajectory regularity (Extended Data Fig. 4a), reflecting the consistency with which animals repeated the reward acquisition cycle—active nose-poke, entering the magazine, receiving the reward, exiting the magazine, and returning to the nose-poke hole. Optimal behavior would correspond to less variation (greater regularity) in their movement patterns^59, 60^. We then performed factor analysis (Extended Data Fig. 4b) to identify a latent factor representing the shared variance across the three behavioral metrics—session time, discrimination ratio, and trajectory regularity. This factor was interpreted as a composite measure of behavioral optimality, reflecting more efficient and accurate performance in the FR1 task.

To validate the robustness of our behavioral optimality factor, and to ensure it was not confounded by general time-dependent trends independent of learning, we performed a leave-one-out factor analysis using residualized behavioral metrics. This validation method involved iteratively hiding each residualized metric (session time, discrimination ratio, trajectory regularity) and correlating it with a latent factor derived from the remaining two residualized metrics (see ‘Validation of behavioral optimality factor using Leave-One-Out factor analysis’ in Methods for details). Crucially, across all three iterations, we observed significant positive repeated-measures correlations^61^ between the hidden residualized behavioral metric and the latent factor (Supplementary Table 3). These consistent and significant positive correlations strongly validate the robustness of our behavioral optimality factor as a reliable measure of task performance, even when accounting for general time-dependent trends and subject variability.

To determine whether behavioral optimality was associated with distinct patterns of cellular Ca^2+^ activity, we analyzed the relationship between the behavioral optimality (factor) and degree of contextual encoding (classification accuracy) for GPe astrocytes and Proto^GPe→STN^ neurons during ME and MX events across both blocks. For GPe astrocytes, we observed no significant correlation between behavioral optimality and classification accuracy for either ME or MX events in the first block (ME: Fig. 3e top, 1^st^ column; MX: Fig. 3f top, 1^st^ column) or the second block (ME: Fig. 3e bottom, 1^st^ column; MX: Fig. 3f bottom, 1^st^ column). In contrast, Proto^GPe→STN^ neurons exhibited a significant positive correlation between behavioral optimality and classification accuracy across both blocks and events (first block, ME: Fig. 3e top, 2^nd^ column; first block, MX: Fig. 3f top, 2^nd^ column; second block, ME: Fig. 3e bottom, 2^nd^ column; second block, MX: Fig. 3f bottom, 2^nd^ column). Furthermore, the correlation coefficients for Proto^GPe→STN^ neurons were significantly higher than those for astrocytes across all cases (First block, ME: Fig. 3e top, 3^rd^ column; First block, MX: Fig. 3f top, 3^rd^ column; Second block, MX: Fig. 3f bottom, 3^rd^ column), except for ME events in the second block, where the difference was not significant (Fig. 3e bottom, 3^rd^ column).

These findings demonstrate that while both Proto^GPe→STN^ neurons and GPe astrocytes exhibit contextual encoding, only the degree of contextual encoding in Proto^GPe→STN^ neurons correlates significantly with behavioral optimality, indicating a tighter link to the adaptive aspects of performance. In contrast, astrocyte-based contextual signals did not vary with behavioral optimality, suggesting they maintain a gradual contextual representation, which could serve a modulatory or maintenance-oriented function in sustaining the overall contextual framework once it is (re-)established, rather than driving rapid adaptation. Given that behavioral optimality in reversal learning depends on behavioral flexibility, our results implicate Proto^GPe→STN^ neurons as plausible contributors to this process through their dynamic and context-dependent activity patterns.

### Chemogenetic inhibition of GPe neurons reduces behavior flexibility

Our previous analyses revealed that Proto^GPe→STN^ neurons exhibit dynamic, context-dependent activity patterns during task learning and reversal phases (Fig. 3d), closely aligning with behavioral adaptation (Fig. 3e-f). Specifically, as animals adapted to new outcome contingencies, Proto^GPe→STN^ neurons dynamically represented context, while astrocyte activity remained gradual. This finding suggests that GPe neurons may directly support behavioral flexibility, enabling animals to modify their actions in response to changing reward contingencies. To test this hypothesis, we employed targeted chemogenetic inhibition to disrupt the elevated activity of GPe neurons immediately following the reversal (Fig. 2d, f), assessing whether this inhibition impairs behavioral flexibility during task reversals.

For the experimental group, we injected a CaMKIIα promoter-driven adeno-associated virus serotype 5 (AAV5) expressing the human M4 muscarinic receptor [hM4D(Gi)] into the GPe. For the control group, we injected a CaMKIIα promoter-driven AAV5 expressing mCherry (Fig. 4a). Two weeks after viral delivery, we confirmed successful viral expression (Extended Data Fig. 5a) and validated the effect of hM4Di activation (Fig. 4c-d and Extended Data Fig. 5b). Treatment with C21, a water-soluble and effective CNO (clozapine N-oxide) analog, significantly reduced the firing rate of GPe neurons (Fig. 4c-d). To evaluate whether chemogenetic inhibition of GPe neurons specifically applied right after a context change affects behavioral flexibility, we administered C21 (3 mg/kg, *i.p.*) 30 minutes before the start of each session following a context reversal – from the initially acquired left context to the new, right context – on Days 10 and 18 (Fig. 4b). This experiment demonstrated that the experimental group exhibited significantly reduced behavioral flexibility during reversal learning (Fig. 4f and Extended Data Fig. 6b) compared to the control group (Fig. 4e and Extended Data Fig. 6a). Specifically, the experimental group showed marked difficulty adapting to the new active nose-poke location during the reversal stages (Fig. 4f and Extended Data Fig. 7), a deficit not observed in the control group (Fig. 4e and Extended Data Fig. 7).

**Figure 4.**
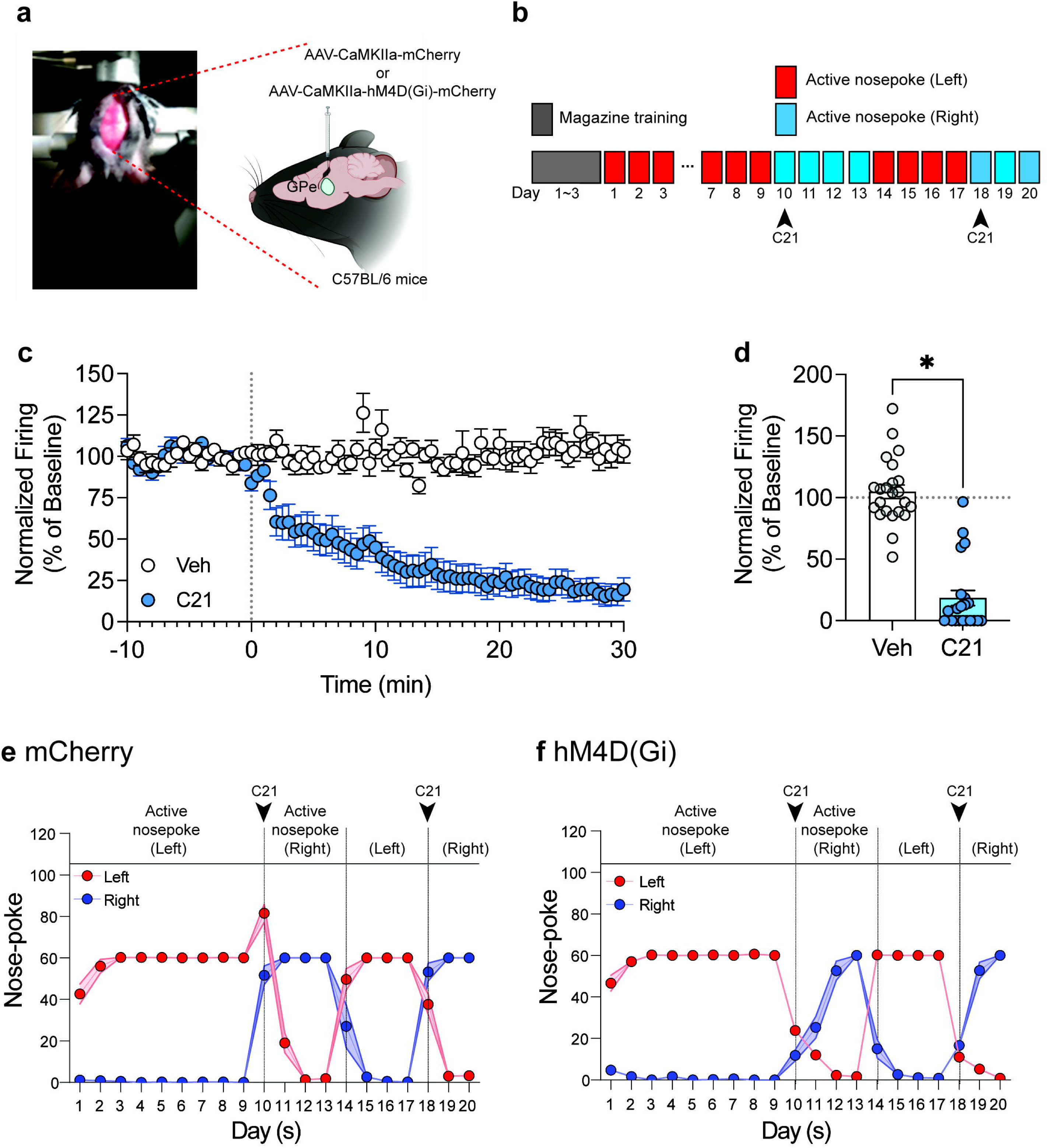
Effect of inactivation of the GPe neurons on behavioral flexibility. **a,** Schematic of viral injections and drug cannula implantations for cell-type-specific expression of inhibitory DREADD [hM4D(Gi)] in GPe neurons. **b,** Schematic of the FR1 task training schedule and drug (compound 21, C21) injection (3 mg/kg, ip). **c,** Time course of normalized firing (percentage with respect to the baseline) following vehicle (Veh) or C21 application (C21). **d,** Pooled data showing significant firing reduction in the C21 group compared to vehicle (\**P* < 0.05, Unpaired t-test). N_Cell_ = 21–23 per group (6 mice). **e-f,** Nose-poking behaviors during the FR1 task training. **e,** Control group (mCherry). **f,** Chemogenetic inhibition group [hM4D(Gi)]. Data are represented as mean ± SEM; Data consist of behavioral performance of 5 mice per group, with identical session counts per block for both groups: *n =* 45 (1^st^ block), 20 (2^nd^ block), 20 (3^rd^ block), and 15 (4^th^ block). See Supplementary Table 1 for full statistical information.

These results indicate that inhibition of GPe neurons impairs reversal learning, highlighting the essential role of GPe neural activity in supporting behavioral flexibility following context changes.

### Caspase-dependent resection of Proto^GPe^**^→^**^STN^ neurons reduces behavior flexibility

To further investigate how GPe neurons contribute to behavioral flexibility, we examined specific GPe projection pathways. The GPe has several projection targets, including the STN, SNr, and lateral habenula (lHb)^25, 26^, each associated with distinct functions in motor control, reward processing, and cognitive flexibility. Given our previous findings of reversal-related activity changes (Fig. 2c-d, f) and contextual encoding (Fig. 3) in GPe prototypic neurons projecting to the STN (Proto^GPe→STN^ neurons), we focused on targeting this pathway. To further confirm the role of Proto^GPe→STN^ neurons in flexible behavior, we used a Cre-dependent caspase 3 system to selectively ablate Proto^GPe→STN^ neurons while minimizing toxicity to neighboring cells^62, 63^. Mice received bilateral injections of a mCherry-tagged retrograde virus expressing Cre recombinase (AAV-Ef1α-mCherry-IRES-Cre7) into the STN, followed by injections of Cre-dependent caspase-3 (AAV-flex-taCasp3-TEVp) into the GPe. Two weeks after caspase-3 administration, mice performed the FR1 task over a longitudinal period to assess their ability to learn and stabilize nose-poke behavior (Fig. 5a).

**Figure 5.**
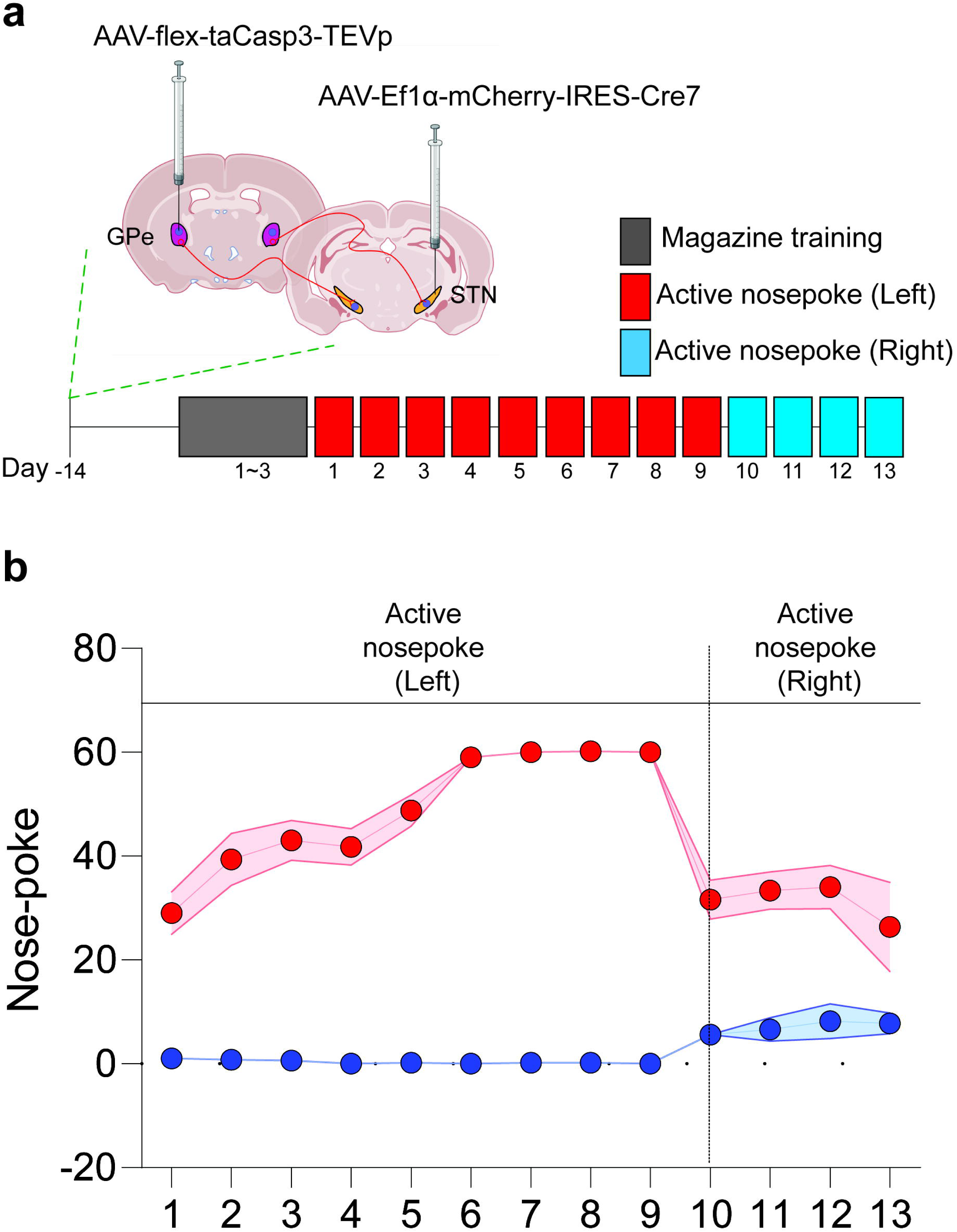
Effect of ablation of the Proto^GPe→STN^ neurons on behavioral flexibility. **a,** Schematic of viral injections for Cre-dependent ablation of Proto^GPe→STN^ neurons and FR1 task training schedule following injection. **b,** Nose-poking behaviors during the FR1 task training. Data are represented as mean ± SEM; from a total of 65 sessions collected across two blocks from 5 mice: *n =* 45 (1^st^ block), and 20 (2^nd^ block). See Supplementary Table 1 for full statistical information.

During the stabilization phase, mice successfully acquired and performed the task without noticeable deficits, indicating that the Proto^GPe→STN^ neurons are not required for the acquisition or stabilization of learned behaviors (Fig. 5b, Days 1-9). However, after the context reversal - where the active nose-poke switched from left to right - mice displayed a marked deficit in behavioral flexibility (Fig. 5b, Day 10). Despite exposure to the new active nose-poke location, they failed to adapt their behavior even four days after the reversal was introduced (Fig. 5b, Days 10-13).

These results validate that Proto^GPe→STN^ neurons are necessary for behavioral flexibility, specifically in enabling adaptation to changing outcome contingencies.

### Activation of DLS iMSNs reduces behavioral flexibility during reversal learning

Given that indirect medium spiny neurons (iMSNs) in the dorsolateral striatum (DLS) are major upstream regulators of Proto^GPe→STN^ neurons ^10, 40^, we tested whether manipulating iMSN activity immediately following a context reversal would affect reversal learning. Following a reversal, Ca^2+^ activity of Proto^GPe→STN^ neurons increased (Fig. 2d, f). If the striatum primarily drives this change (Fig. 6a left), then activating iMSNs after the reversal would disrupt reversal learning by decreasing Proto^GPe→STN^ neuronal activity, thereby impairing behavioral flexibility (Fig. 6a left, 2^nd^ row of the table). Conversely, we predicted that inhibiting iMSNs post-reversal would not affect behavioral flexibility, as this aligns with the natural mechanism triggered by context change – reduced iMSN activity leading to increased Proto^GPe→STN^ neuronal activity (Fig. 6a left, 3^rd^ row of the table).

**Figure 6.**
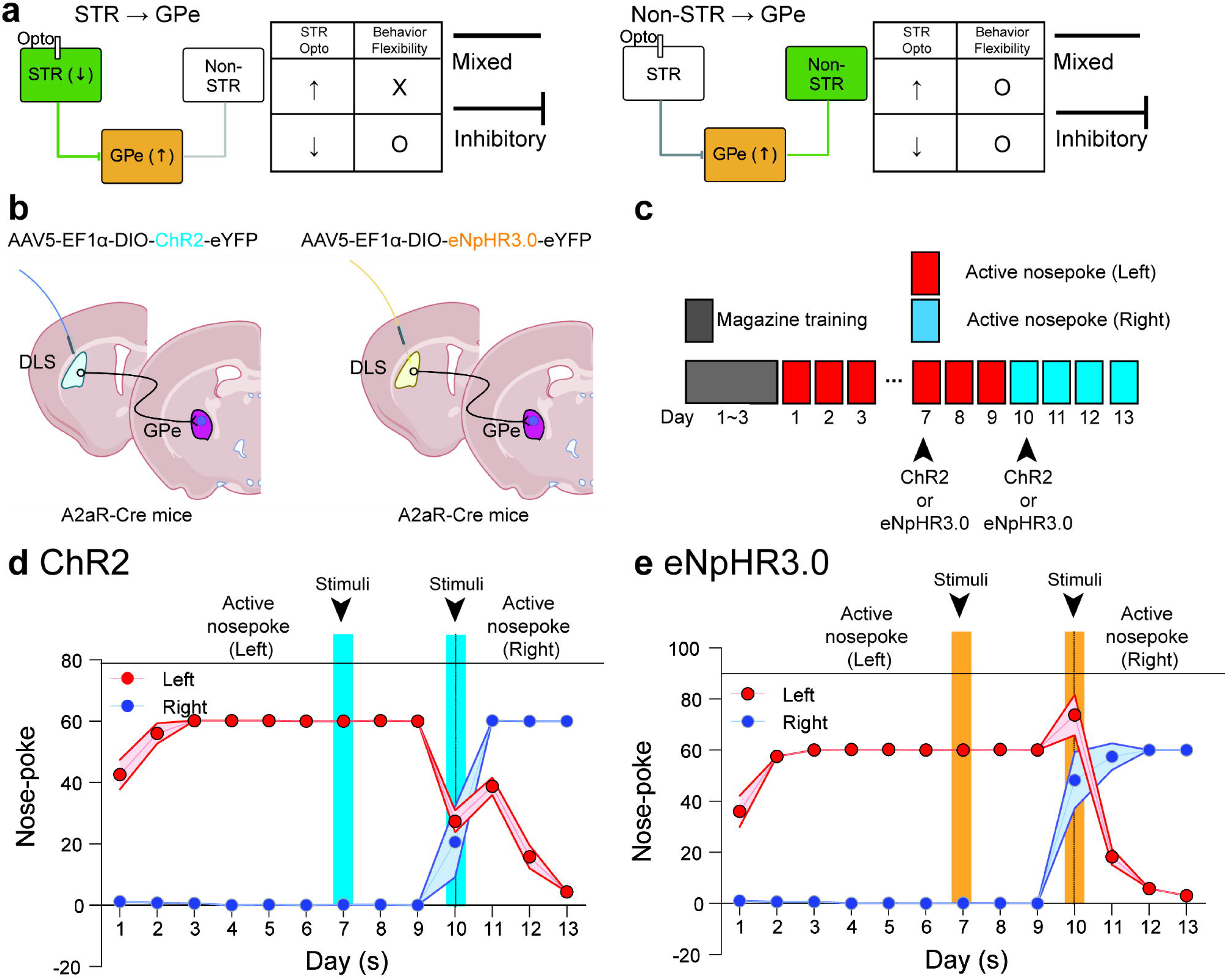
Optogenetic manipulation of DLS iMSNs alters behavioral flexibility. a, Schematic depicting two hypothetical mechanisms for input modulating the Proto^GPe→STN^ neurons. Inhibitory projections are depicted as flat-ended arrows, and non- striatal inputs (mixed; excitatory and inhibitory) as lines without arrowheads. In each mechanism, green highlights the primary region/pathway, while grey indicates non- primary inputs/regions. Left: DLS iMSNs as the primary input driving Proto^GPe→STN^ neurons. Right: Non-striatal inputs primarily drive Proto^GPe→STN^ neurons. **b,** Schematic of viral injections and optic fiber implantations for optogenetic activation (left, ChR2) and inhibition (right, eNpHR3.0) of DLS iMSNs. **c,** Schematic of the FR1 task training schedule and laser delivery for optogenetic manipulation. **d-e,** Nose-poking behaviors during the FR1 task training. **d,** Optogenetic activation group (ChR2), with optogenetic stimulation on days 7 and 11-12 indicated an increase in nose-poke behavior in response to stimulation. **e,** Optogenetic inhibition group (eNpHR3.0). No change in nose-poke behavior when stimulated on day 7 and day 11-12. Data are represented as mean ± SEM. Data consist of behavioral performance of 5 mice per group, with identical session counts per block for both groups: *n =* 45 (1^st^ block), and 20 (2^nd^ block). STR, striatum; GPe, external globus pallidus; Non-STR, non-striatal inputs.See Supplementary Table 1 for full statistical information.

Alternatively, Proto^GPe→STN^ neurons also receive input from various other regions, including the STN, cortex, thalamus, and internal globus pallidus (GPi). If the regulation of Proto^GPe→STN^ neurons in behavioral flexibility is predominantly driven by these other regions (Fig. 6a right), then manipulating iMSN activity would not be expected to influence behavioral flexibility (Fig. 6a right, table).

To test these conflicting mechanisms, we examined whether optogenetic manipulation of DLS iMSNs influences behavioral flexibility during longitudinal FR1 tasks, with a focus on reversal learning. To assess the role of DLS iMSNs in reward-seeking and adaptive behavior, we injected a Cre-dependent ChR2-expressing virus into the DLS of A2aR-Cre mice and implanted optical fibers in the DLS (Fig. 6b-c). This setup enabled selective activation of DLS iMSNs through optogenetic stimulation (Extended Data Fig. 8). Optogenetic activation of DLS iMSNs using ChR2 significantly impaired animals’ ability to adapt during reversal learning (Fig. 6d, Day 10). Mice with activated DLS iMSNs exhibited reduced behavioral flexibility, failing to switch to the newly active nose-poke following the reward contingency reversal. Conversely, optogenetic inhibition of DLS iMSNs using eNpHR3.0 did not affect reversal learning behavior (Fig. 6e, Day 10). Mice subjected to DLS iMSN inhibition displayed behavioral flexibility comparable to control animals.

To determine whether the observed deficits in reversal learning were attributable to changes in movement speed^39, 44^ or punishment effects^44^ associated with iMSN manipulation, we performed optogenetic manipulation during the first block, when mice had successfully acquired the task strategy (Fig. 6d-e, Day 7). We compared session times and discrimination ratios on the manipulation day (Day 7) with the days immediately before (Day 6) and after manipulation one day after manipulation (Day 8). Our analysis revealed no significant differences in session times or discrimination ratios for the optogenetic activation group (Extended Data Fig. 9a and Fig. 6d). Similarly, no significant changes in session times or discrimination ratios were observed for the optogenetic inhibition group (Extended Data Fig. 9b and Fig. 6e). These results indicate that the deficits observed in reversal learning (Fig. 6d) cannot be attributed to changes in movement speed or punishment effects.

These findings suggest that DLS iMSNs provide a modulatory input to Proto^GPe→STN^ neurons (Fig. 6a left). When reward contingencies change, Proto^GPe→STN^ neurons incorporate reduced input from DLS iMSN and utilize it to support behavioral flexibility.

### STN excitatory input to Proto^GPe^**^→^**^STN^ neurons underpins behavioral flexibility

Building upon our findings implicating Proto^GPe→STN^ neurons and their striatal inputs in behavioral flexibility, we next investigated the role of STN, another major afferent to these GPe neurons^10, 64^. The canonical basal ganglia indirect pathway model^27, 28^ (Fig. 7a left) traditionally describes basal ganglia information flow as Striatum→GPe→STN→SNr, where the STN receives inhibitory input from the GPe (GPe→STN) and then modulates downstream circuits, such as the SNr. Within this ‘canonical information flow’ framework, our observation of increased Proto^GPe→STN^ neuron activity after reversal (Fig. 2d, f) predicts a decrease in STN activity, ultimately reducing STN output to the SNr. Therefore, canonical information flow theory predicts: activating STN neurons after reversal should disrupt reversal learning (Fig. 7a left, 3^rd^ row of the table), since it counteracts with the inhibitory input given by the GPe. Conversely, inhibiting STN neurons would align with ‘canonical information flow’ and not affect behavioral flexibility (Fig. 7a left, 2^nd^ row of the table).

**Figure 7.**
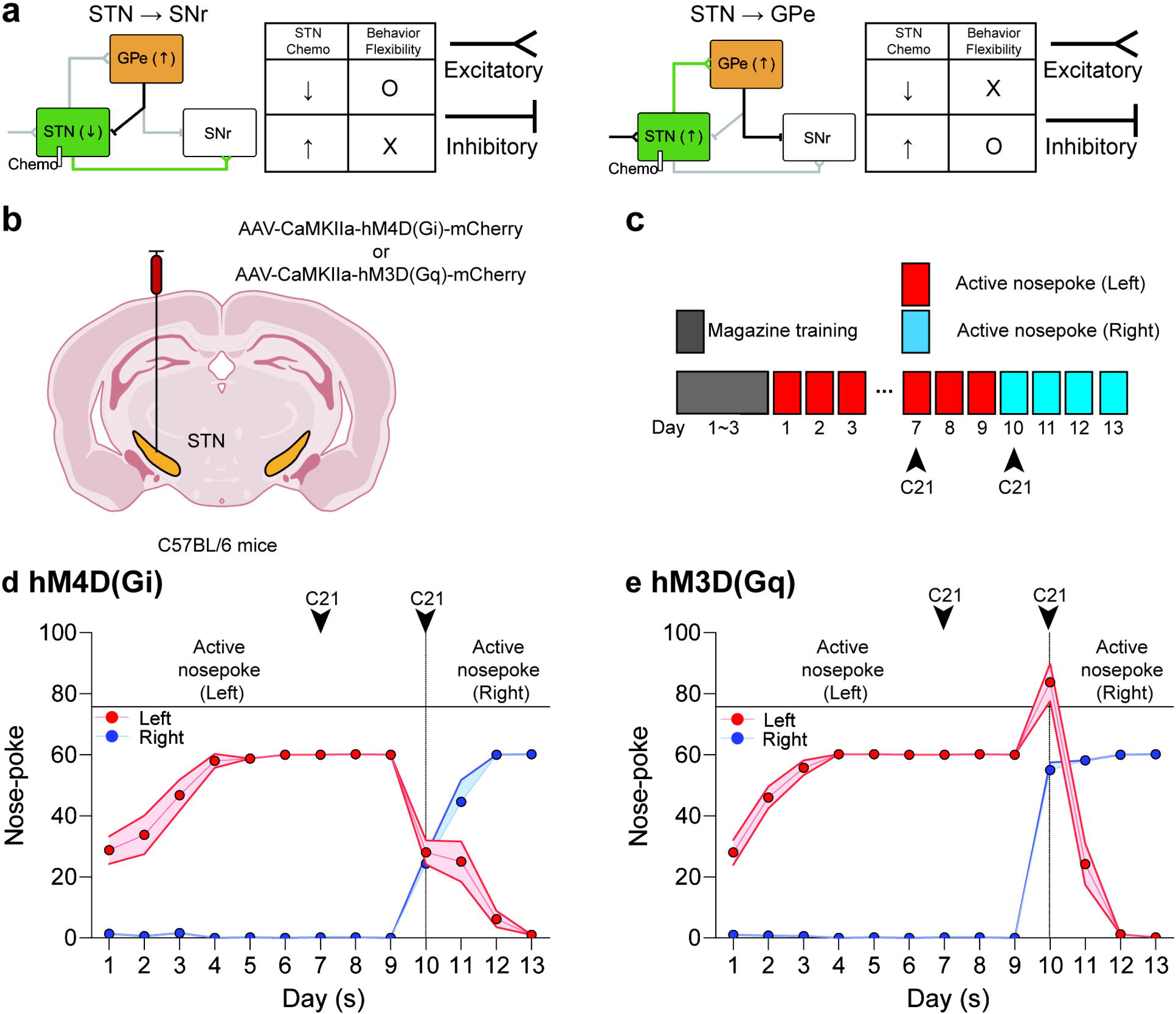
Chemogenetic manipulation of STN neurons alters behavioral flexibility. **a,** Schematic showing two hypothetical roles of the STN in the Proto^GPe→STN^ neuron- mediating circuit. Excitatory projections are shown as arrows with triangular tips, and inhibitory projections are flat-ended arrows. In each mechanism, green highlights the primary region/pathway, while grey indicates non-primary inputs/regions. Black projections represent pathways that drive or relay information according to each hypothesis. Left: STN as a relay nucleus (canonical information flow model). Right: STN as an input (non-canonical information flow model). **b,** Schematic of viral injections and drug cannula implantations for cell-type-specific expression of inhibitory [hM4D(Gi)] or excitatory [hM3D(Gq)] DREADD in STN neurons. **c,** Schematic of the FR1 task training schedule and drug (compound 21, C21) injection (3 mg/kg, ip). **d-e,** Nose-poking behaviors during the FR1 task training. **d,** Chemogenetic inhibition group [hM4D(Gi)]. Chemogenetic inhibition (hM4D(Gi)) of STN neurons reduces nose-poke behavior after C21 administration on days 7 and 11-12. **e,** Chemogenetic activation group [hM3D(Gq)]. Chemogenetic activation of STN neurons (hM3D(Gq)) did not alter nose-picking behavior after C21 administration on days 7 and 11-12. Data are represented as mean ± SEM. Data consist of behavioral performance of 5 mice per group, with identical session counts per block for both groups: *n =* 45 (1^st^ block), and 20 (2^nd^ block). GPe, external globus pallidus; STN, subthalamic nucleus; SNr, substantia nigra pars reticulata. See Supplementary Table 1 for full statistical information.

However, substantial recent evidence suggests a more complex reality, challenging this simplified ‘canonical’ picture of basal ganglia information flow. This includes a direct projection of Proto^GPe→STN^ neurons to the SNr via axonal collateral^10, 34^, a hyperdirect pathway from cortex to STN^35–38^, and a feedback circuit from the STN to the GPe^10, 39, 40^. These findings support an alternative, ‘non-canonical’ information flow’ model (Fig. 7a right), where Proto^GPe→STN^ neurons receive excitatory input from the STN (STN→GPe) and, in turn, modulate downstream circuits. In this ‘non-canonical information flow’ scenario, increased Proto^GPe→STN^ neuron activity arises from enhanced STN activity. Consequently, ‘non-canonical information flow’ predicts the opposite outcome: inhibiting the STN neurons right after the reversal should impair reversal learning (Fig. 7a right, 2^nd^ row of the table) by blocking the natural rise in Proto^GPe→STN^ neuron activity, which is necessary for Proto^GPe→STN^ neurons to control the downstream circuit. Conversely, activating STN neurons would align with the ‘non-canonical information flow’ and leave flexibility intact (Fig. 7a right, 3^rd^ row of the table).

To test these mechanisms, we microinjected AAV-CaMKIIα-hM4D(Gi)-mCherry or AAV-CaMKIIα-hM3D(Gq)-mCherry into the STN of C57BL/6 mice (Fig. 7b), enabling selective chemogenetic inhibition or activation of STN neurons. After confirming viral expression, we administered C21 (3 mg/kg, *i.p.*) to either inhibit or activate STN neurons and assessed the animals’ performance during a reversal learning task (Fig. 5g-h). Chemogenetic inhibition of STN neurons using hM4D(Gi) significantly impaired behavioral flexibility (Fig. 7d, Day 10). Conversely, chemogenetic activation of STN neurons with hM3D(Gq) did not affect behavioral flexibility (Fig. 7e, Day 10). To confirm that the observed deficits in reversal learning were not attributable to changes in movement speed^38^ or aversive learning^65^ potentially associated with STN manipulation, we performed chemogenetic manipulations during the first block when mice had already acquired the necessary task strategy (Fig. 7d-e, Day 7). We analyzed session times and discrimination ratios on the manipulation day (Day 7) and compared them to the day before (Day 6) and the day after the manipulation (Day 8). The results revealed no significant differences in session time (Extended Data Fig. 9d) for the chemogenetic activation group. Similarly, no significant differences were observed in session time (Extended Data Fig. 9c) for the chemogenetic inhibition group. These results show that the observed deficits in reversal learning were not attributable to altered movement speed^38^ or aversive learning^65^ effects but were specifically related to the manipulation of STN neurons.

These findings demonstrate that Proto^GPe→STN^ neurons require excitatory input from STN neurons to maintain behavioral flexibility during reversal learning. When reward contingencies change, Proto^GPe→STN^ neurons incorporate increased input from the STN and utilize it to support behavioral flexibility.

## Discussion

Here, we identify the necessity of Proto^GPe→STN^ neurons in facilitating behavioral flexibility. Using a longitudinal operant conditioning task with context reversals, we show that Proto^GPe→STN^ neurons develop dynamic contextual encoding that correlates with behavioral optimality, underscoring their importance in adapting to changing environmental contingencies. Critically, the deletion of Proto^GPe→STN^ neurons impairs the ability to adjust behavior following context reversals, yet does not affect initial reward-seeking acquisition, directly demonstrating the pathway’s specific role in enabling behavioral flexibility. Chemogenetic and optogenetic manipulations of DLS iMSNs and STN neurons revealed that Proto^GPe→STN^ neurons integrate inputs from both regions to modulate downstream basal ganglia activity via direct projections to the SNr via axonal collateral (GPe→SNr)^10, 34^, rather than through STN mediation (GPe→STN→SNr) (Fig. 8).

**Figure 8.**
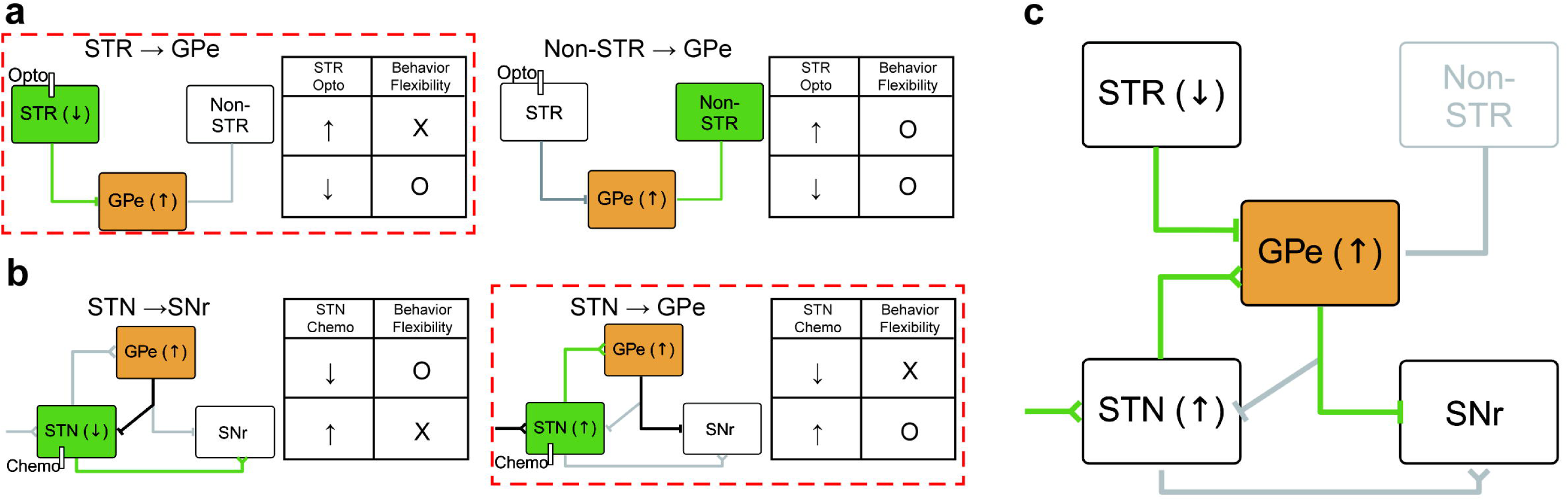
Schematics summarizing the results of the present study. Excitatory projections are shown as arrows with triangular tips, inhibitory projections as flat-ended arrows, and non-striatal inputs (mixed; excitatory and inhibitory) as lines without arrowheads. The mechanism that aligns with experiment data is enclosed within the red dotted rectangle. **a,** Two hypothetical mechanisms for input modulating the Proto^GPe→STN^ neurons (Fig. 6). Left: DLS iMSNs as the primary input driving Proto^GPe→STN^ neurons. Right: Non-striatal inputs primarily drive Proto^GPe→STN^ neurons. **b,** Two hypothetical roles of the STN in the Proto^GPe→STN^ neuron-mediating circuit (Fig. 7). Left: STN as a relay nucleus (canonical information flow model). Right: STN as an input (non-canonical information flow model). **c,** Summary diagram of the study’s findings. Proto^GPe→STN^ neurons enable behavioral flexibility by integrating striatal and subthalamic inputs and sending output directly to the SNr, without STN mediation. Green lines represent the major pathways driving this mechanism, while gray lines indicate secondary pathways. STR, striatum; GPe, external globus pallidus; STN, subthalamic nucleus; SNr, substantia nigra pars reticulata; Non-STR, non-striatal inputs.

Our findings concur with the ’non-canonical information flow’ model of basal ganglia circuitry for behavioral flexibility, revealing a critical and previously underappreciated role for Proto^GPe→STN^ neurons as integrators of excitatory STN input. Specifically, our chemogenetic manipulation of STN neurons directly tested the contrasting predictions of the ’canonical’ versus ’non-canonical information flow’ models. The observation that inhibition of STN neurons impaired reversal learning, while activation did not, demonstrates that Proto^GPe→STN^ neurons require excitatory drive from the STN to enable adaptive behavior changes. This finding is further corroborated by our optogenetic manipulation of DLS iMSNs, showing that reduced inhibitory input from the striatum is also critical for Proto^GPe→STN^ neuron function in flexibility. Taken together, our manipulations of both STN and DLS inputs converge to highlight the ’non-canonical information flow’ model, where Proto^GPe→STN^ neurons actively integrate convergent excitatory and inhibitory signals to orchestrate flexible behavior.

Furthermore, our findings demonstrate that Proto^GPe→STN^ neurons are essential for behavioral flexibility. Caspase-dependent ablation of Proto^GPe→STN^ neurons revealed that initial reward-seeking acquisition was unaffected, but adaptive responses to changing action-outcome contingencies were significantly impaired, directly demonstrating their specific role in enabling behavioral flexibility. This contrasts with a previous study that suggested Proto^GPe→Pf^ neurons, not Proto^GPe→STN^ neurons, are partially involved in reversal learning^10^. Specifically, they showed that optogenetic activation of Proto^GPe→Pf^ axons, but not Proto^GPe→SNr^ axons (neurons that also project to the STN via axonal collaterals^10, 34^), impaired a foraging task with context reversal. Critically, they also reported that manipulation of neither Proto^GPe→Pf^ nor Proto^GPe→SNr^ neurons (Proto^GPe→STN^ neurons) affected performance in an operant conditioning task with context reversal – the same task we employed in our study.

The apparent discrepancy may be attributed to the differences in the phase of learning during which the reversal occurs, as well as the distinct neural mechanisms engaged at these stages. In this study^10^, context reversals were introduced at a time when subjects exhibited relatively low accuracy (∼80%), suggesting that continued learning could further enhance performance. Also, subjects had limited exposure to the experimental environment (< 90 min)^10^, characterizing this as the early phase of acquisition. These differences likely reflect the dynamic recruitment of GPe circuits as learning progresses, highlighting phase-specific roles for Proto^GPe→STN^ and Proto^GPe→Pf^ neurons in behavioral flexibility. Of note, our study implemented reversals after animals had already achieved high accuracy (∼100%, Extended Data Fig. 1) and fully acclimated to the environment (∼240 min, Extended Data Fig. 1), representing a more stable, late phase of acquisition. Previous studies have highlighted a transition in behavioral control as training or learning progresses, shifting from dorsomedial striatum (DMS)-mediated to dorsolateral striatum (DLS)-mediated regulation^66, 67^. Within the GPe, Proto^GPe→Pf^ neurons predominantly receive input from the DMS, whereas Proto^GPe→STN^ neurons are primarily targeted by the DLS^10^. This anatomical distinction suggests that the recruitment of specific GPe circuits depends on which striatal region exerts control during different training phases. During the early phase of acquisition, the DMS-driven Proto^GPe→Pf^ circuit likely governs action selection and behavioral flexibility. As learning advances to the late phase, with control shifting to the DLS, the Proto^GPe→STN^ circuit becomes the dominant contributor, reflecting the stronger DLS input at this phase. Therefore, when a context reversal occurs in the late phase, the Proto^GPe→STN^ circuit mediates behavioral flexibility. This phase-dependent recruitment underscores the dynamic engagement of distinct GPe circuits during training and adaptation. Further investigation into the transition of control from Proto^GPe→Pf^ neurons to Proto^GPe→STN^ neurons could elucidate how the GPe supports behavioral flexibility across different phases of learning, advancing our understanding of basal ganglia circuit dynamics.

Unlike Proto^GPe→STN^ neurons, GPe astrocytes exhibited distinct patterns of contextual encoding. Contextual encoding of astrocytes remained stable during the initial acquisition and stabilization of behavioral strategies and was unaffected by context reversals. However, following a reversal, astrocytes demonstrated an increasing degree of contextual encoding as learning progressed. These observations align with the contextual guidance model^68^, which proposes that astrocytes might adjust their responses according to contextual cues. According to this framework, astrocytes dynamically reconfigure neural networks in response to contextual signals, thereby modulating behavior. During the initial acquisition phase, when no context changes occur, astrocytes exhibit stable activity patterns maintaining a consistent baseline, manifesting as a constant degree of contextual encoding during the first block (Fig. 3d, GFAP+, Days 1-29). This stability may enable the neural network, including Proto^GPe→STN^ neurons, to refine its activity patterns, facilitating behavioral optimization. This refinement is evidenced by an increasing degree of contextual encoding in Proto^GPe→STN^ neurons during the first block, which correlates strongly with behavioral optimality (Fig. 3d, Proto, Days 1-29).

Immediately following a context reversal, astrocytes can receive modulatory signals conveying information about the change in context, such as neuromodulators, sensory inputs, or neurohormones, reflecting the animal’s internal and external state^69,70^. In response, astrocytes begin altering their activity when reward contingencies change. Compared to rapid point-to-point neuronal transmission, the slower kinetics of these modulatory signals may drive more consistent and gradual changes in astrocyte activity^71^, aligning with the delayed but sustained rise in contextual encoding we observed (Fig. 3d, GFAP+, Days 30-46). As a result, astrocytes develop distinct activity patterns compared to those observed during the initial acquisition phase, as evidenced by an increasing degree of contextual encoding in astrocytes during the second block compared to the first block (Fig. 3c, red violins, LNP versus RNP). Although the precise mechanisms remain to be tested, it is plausible that these changes in astrocytic activity influence local network dynamics^47, 72^. This change could then trigger a temporary decrease in the contextual encoding of Proto^GPe→STN^ neurons right after the reversal (Fig. 3d, Proto, Day 30). As the network continues to adapt and learn the new context, Proto^GPe→STN^ neurons refine their activity patterns, ultimately leading to more optimal behavior. This is reflected in an increasing degree of contextual encoding during the second block, correlating strongly with behavioral optimality (Fig. 3d, Proto, Days 30-46).

Collectively, under the contextual guidance model^68^, our results suggest that GPe astrocytes may support behavioral flexibility by maintaining a background or consolidated representation of the current context, potentially contributing to the overall stability or reliability of GPe network activity. Given our recent study showing that GPe astrocytic dynamics regulate the transition from goal-directed to habitual reward-seeking behavior^55^ —a process intricately linked to behavioral flexibility—further exploration of the role of GPe astrocytes is warranted. Such investigations could provide critical insights into how the basal ganglia circuit supports adaptive behavior, elucidating the contributions of astrocytic activity to the modulation of neural network dynamics during behavioral transitions.

While our findings highlight the role of Proto^GPe→STN^ neurons and GPe astrocytes in behavioral flexibility, it is important to acknowledge the cellular diversity within GPe. Besides Proto^GPe→STN^ neurons and GPe astrocytes, the GPe contains distinct populations such as arkypallidal neurons and non-parvalbumin-expressing subpopulations, each contributing uniquely to basal ganglia circuitry and the regulation of motor and cognitive processes^26, 42, 72, 73^. For instance, arkypallidal neurons, which predominantly project back to the dorsal striatum, have been implicated in the regulation of habitual seeking behaviors^63^, suggesting a potential role in behavioral flexibility. Furthermore, choline acetyltransferase-expressing (ChAT+) neurons, which project to cortical regions^74^, such as the orbitofrontal^75^ and medial prefrontal cortex^76^, may also influence behavioral adaptation given the involvement of these cortical areas in flexible decision making^3–5^. These diverse cellular contributions underscore the multifaceted role of GPe subpopulations in shaping adaptive behavior and highlight the need for further investigation into their specific roles within basal ganglia networks.

Local interactions within the GPe are critical for shaping its output and functional dynamics. Proto^GPe→STN^ neurons provide robust inhibitory input to neighboring neurons through reciprocal connections, thereby influencing local circuit activity^32, 34, 39, 40^. Additionally, astrocytes as the most abundant cell type in the GPe^45^, play an integral role in modulating nearby neural circuits^51–53^, further enhancing the capacity of the GPe to fine-tune basal ganglia activity in response to dynamic environmental demands. These local interactions form a complex network that supports the adaptability required for behavioral flexibility. Future studies exploring the diversity of GPe cell types and their local interactions will be essential for constructing a more comprehensive blueprint of the GPe’s contribution to adaptive behavior.

To delineate the functional circuit responsible for behavioral flexibility, we first validated the necessity of Proto^GPe→STN^ neurons through caspase-dependent pathway resection. To further understand which inputs Proto^GPe→STN^ neurons integrate and the output pathways they utilize for behavioral flexibility, we tested conflicting mechanisms by making predictions about the effects of manipulating specific regions of interest and comparing these predictions to the outcomes of manipulation experiments. However, it is important to note that these predictions are based primarily on the direct interactions between the region of interest and the GPe.

The basal ganglia operate as a highly interconnected network, where regions influence one another both directly and indirectly^77^. For instance, while the DLS provides direct inhibitory input to the GPe and the STN provides direct excitatory input, their effects on GPe activity extend beyond these singular pathways. The DLS and STN also exert influence through multi-step connections mediated by intermediary structures, including dopaminergic brainstem nuclei^78, 79^, thalamic intralaminar nuclei^80, 81^, and cortical regions^75, 82, 83^. These indirect pathways form effective connections between the DLS, STN, and GPe, underscoring the importance of considering broader network dynamics when interpreting striatal and subthalamic contributions to GPe function. Future studies should employ computational models that integrate diverse projections and interactions across basal ganglia nuclei to capture the complexity of these interactions. Such models will be instrumental in unraveling the dynamic, multifaceted contributions of these circuits to behavioral flexibility.

In summary, our study identifies Proto^GPe→STN^ neurons as an important node within the basal ganglia circuitry, integrating striatal and subthalamic inputs to dynamically regulate adaptive behavior. These findings reveal a non-canonical role for Proto^GPe→STN^ neurons, extending beyond traditional relay functions to establish them as pivotal modulators supporting context-driven behavioral flexibility, operating within a ’non-canonical information flow’ architecture of the basal ganglia. Understanding this specialized function of Proto^GPe→STN^ neurons opens new avenues for investigating and potentially treating disorders characterized by impaired behavioral adaptation^1, 2^, such as obsessive-compulsive disorder, autism spectrum disorder, Parkinson’s disease, and addiction, highlighting the broader translational relevance of dissecting specific GPe circuit mechanisms.

## Methods

### Animals

All experimental procedures were conducted with guidelines set by the Mayo Clinic Institutional Animal Care and Use Committee, adhering to the principles outlined by the National Institutes of Health. We used C57BL/6J mice (stock no. 000664) mice and GFAP-Cre (stock no. 024098)/DIO-GCaMP6s (stock no. 028866) bi-transgenic mice (Jackson Laboratory, Bar Harbor, ME). The animals were housed in transparent Plexiglas cages, with environmental conditions maintained at a stable temperature of 24°C ± 1°C and humidity levels of 60 ± 2%. The room followed a 12-hour light/12-hour dark cycle, with lights turning on at 7:00 a.m. All experiments were performed on 8- to 10-week-old male mice. Mice had unrestricted access to food and water, except during operant conditioning tasks, during which they were food-restricted to maintain 85% of their original weight throughout the experiment.

### Stereotaxic surgery and virus injection

Mice were anesthetized with isoflurane (1.5% in oxygen gas) with the VetFlo vaporizer with a single-channel anesthesia stand (Kent Scientific Corporation, Torrington, CT) and placed on the digital stereotaxic alignment system (model 1900, David Kopf Instruments, Tujunga, CA). Hair was trimmed, and the skull was exposed using an 8-gauge electrosurgical skin cutter (KLS Martin, Jacksonville, FL). The skull was leveled using a dual tilt measurement tool. Holes were drilled in the skull at the appropriate stereotaxic coordinates. For chemogenetics, viruses were infused unilaterally to the GPe [anteroposterior (AP) −0.46 mm, mediolateral (ML) +2.0 mm, dorsoventral (DV) −3.2 mm from dura] at 100 nl/min for 3 min through a 33-gauge injection needle (catalog no. NF33BV, World Precision Instruments) using a micro syringe pump (model UMP3, World Precision Instruments). The injection needle remained in place for an additional 6 min following the end of the injection. We injected buprenorphine sustained-release LAB (1 mg/kg, subcutaneously; ZooPharm, Laramie, WY) to alleviate post-surgery pain. Mice were used for the experiments 4 to 5 weeks after the virus injection. We injected viruses at the following titers: AAV5-GFAP-mCherry (for control), 1.6×10^13^ GC/ml (Vector Biolabs, Malvern, PA); AAV5-GFAP-hM3Dq-mCherry, 1.2×10^13^ GC/ml. AAV-GFAP-hM3Dq-mCherry was a gift from B. Roth [Addgene viral prep no. 50478-AAV5; http://n2t.net/addgene:50478; Research Resource Identifiers (RRID): Addgene_50478]. AAV5-CamKIIα-hM4Di-mCherry, 2×10^12^ GC/ml (Addgene viral prep no. 50477), AAV5- Ef1a-DIO hChR2-eYFP, 1×10¹³ GC/ml (Addgene viral prep no. 35509-AAV5), AAV5- Ef1a-DIO eNpHR 3.0-eYFP, 1×10¹³ GC/ml (Addgene viral prep no. 26966-AAV5).

### Immunofluorescence

Brains were fixed with 4% paraformaldehyde (Sigma-Aldrich) and transferred to 30% sucrose (Sigma-Aldrich) in phosphate-buffered saline at 4°C for 72 hours. Brains were then frozen in dry ice and sectioned at 40 μm using a microtome (Leica Corp., Bannockburn, IL). Brain slices were stored at −20°C in a cryoprotectant solution containing 30% sucrose (Sigma-Aldrich) and 30% ethylene glycol (Sigma-Aldrich) in phosphate-buffered saline. Sections were incubated in 0.2% Triton X-100 (Sigma-Aldrich) and 5% bovine serum albumin in phosphate-buffered saline for 1 hour, followed by incubation with the primary antibody in 5% bovine serum albumin overnight at 4°C. The Primary antibodies used in the present study included mouse anti-GFAP Alexa Fluor 488 (1:100; monoclonal immunoglobulin G1, #53-9892-82, Thermo Fisher Scientific, Waltham, MA) antibody. After washing with phosphate-buffered saline three times, the sections were mounted onto a glass slide coated with gelatin and cover- slipped with a VECTASHIELD antifade mounting medium with 4′,6-diamidino-2- phenylindole (Vector Laboratories, Burlingame, CA). Images were obtained using an LSM 700 laser scanning confocal microscope (Carl Zeiss, Heidelberg, Germany) using a 10× lens (Extended Data Fig. 4) and a 63× water-immersion lens.

### Chemogenetics

Compound 21 (C21) was obtained from Hello Bio (Princeton, NJ). Following established protocols, mice were administered C21 (3 mg/kg). Previous studies have shown that these doses modify reward-seeking behaviors with minimal impact on motor function or seizure risk^55, 84–86^. In the chemogenetic experiments, operant conditioning with 20% sucrose was first conducted in an FR1 schedule using two cohorts of mice (mCherry *n* = 10, hM4Di *n* =10).

### Optogenetics

During the FR1 experiment, unilateral optogenetic stimulation of channelrhodopsin-2 (ChR2) was performed using a 473 nm laser, and unilateral optogenetic inhibition of enhanced halorhodopsin (eNpHR) was performed using a 593 nm laser. The light was delivered using a 2 ms pulse width, 10 Hz, and 3 mW of power measured at the end of a 200 m optical fiber, while mice were free to move in the operant conditioning chamber for up to 60 min.

### *In vivo* calcium imaging with fiber-photometry

We monitored real-time cellular calcium transients *in vivo* using fiber photometry. An optic cannula (200/240 μm diameter, 200 μm end fiber) was surgically implanted into the GPe of GFAP-Cre/DIO-GCaMP6s mice (coordinates: AP −0.46 mm, ML +2.0 mm, DV −3.0 mm from the dura). The cannula was connected to a patch cord, maintaining a light intensity of 60 μW at the fiber tip.

We used the Multi-Wavelength Photometry System (version 1.2.0.14, Plexon) with time-division multiplexing to acquire signals from multiple wavelengths simultaneously. We used 410 nm as an isosbestic control to correct for calcium-independent fluorescence, movement artifacts, and photobleaching, 465 nm for the excitation of GCaMP6s (astrocyte calcium indicator), and 560 nm for the excitation of jRGECO1a (Proto^GPe→STN^ neuron calcium indicator). A custom MATLAB script calculated the relative fluorescence change (ΔF/F). To correct for calcium-independent fluctuations, the 410 nm control signal was linearly fitted to the 465 nm signal (astrocytes) and separately to the 560 nm signal (Proto^GPe→STN^ neurons). The fitted 410 nm signal was then used to normalize each corresponding signal as follows: ΔF/F = [465 nm signal − fitted 410 nm signal]/fitted 410 nm signal for astrocytes, and ΔF/F = [560 nm signal − fitted 410 nm signal]/fitted 410 nm signal for Proto^GPe→STN^ neurons.

For z-scored normalization, ΔF/F data from GPe astrocytes and Proto^GPe→STN^ neurons were processed separately. For each subject, ΔF/F data across 54 days were collected, and the mean and standard deviation were calculated from this dataset. The calculated mean and standard deviation for each subject were then used to z-score the ΔF/F data from individual sessions of that same subject, ensuring normalization was specific to each subject. The camera capturing the mice’s behavior was synchronized with the fiber photometry recordings, recorded at 30 frames per second. Calcium signals in GPe were recorded daily for 54 days under operant conditioning.

### *In vivo* electrophysiology

The experiments were performed as described previously^54, 55^. Briefly, mice, 3 to 4 weeks following virus injections, were anesthetized by intraperitoneal injection of urethane (1.5 g/kg; Sigma-Aldrich, St. Louis, MO) and placed horizontally on a stereotaxic frame (RWD Life Science, San Diego, CA). We constantly monitored respiratory rate and pedal withdrawal reflex during anesthesia, and the physiological body temperature was maintained using a small-animal feedback-controlled warming pad (RWD Life Science, San Diego, CA). After the scalp incision, small burr holes were drilled to insert high-impedance microelectrodes (Cambridge NeuroTech, Cambridge, UK). We placed the reference wire (Ag/AgCl, 0.03 inches in diameter, A-M systems) in the contralateral parietal cortex. Electrophysiological signals were digitized at 20 kHz and bandpass–filtered from 300 to 5000 Hz (RHD Recording system, Intan Technologies, Los Angeles, CA). The data were analyzed with Clampfit (version 11.2, Molecular Devices, San Jose, CA) and a custom-written code in MATLAB (R2019a, The MathWorks, Natick, MA). For the recording combined with optogenetics to evaluate the circuit from DLS to GPe, a blue LED (Peak wavelength range: 450-465 nm, Optogenetics-LED-STSI module, Prizmatix) was used to deliver unilateral optogenetic stimulation of ChR2-eYFP. The light was delivered using 5-ms pulse widths, 20Hz, and ∼0.6 mW of power measured at combined end faces of both 200 µm optical fibers (Prizmatix).

### Behavioral experiments (Operant conditioning)

The operant chamber consisted of an active hole, an inactive hole, a magazine, a house light, a speaker, and a cue light at each nose port. Reward (20% [w/v] sucrose solution) was presented to the liquid receptacle in the magazine once per reward signal by a syringe pump. Rewards are determined by nose-poking location. When mice performed a nose-poke in the active hole (rewarded or active nose-poke), the chamber presented a tone, light from the nose port, and one reward from the magazine (Fig. 1b). Conversely, no cues or rewards were presented when the nose-poke was performed in the inactive hole (non-rewarded or inactive nose-poke) (Fig. 1b). The shared structure of operant conditioning schedule is as follows (Fig. 1a, 4b, 5a, 6c, 7c).

On days one to three, magazine training was conducted for mice to learn to obtain 60 rewards in the magazine’s space. Following this, the fixed ratio of 1 (FR1) was performed with a maximum session time of 60 minutes. If mice obtained 60 rewards, the session was terminated regardless of the remaining time. For rapid learning of operant behavior, 10 ul of 20% sucrose was placed in the active hole as bait before starting the FR1 session. Moreover, in two consecutive sessions, when the average latency time from nose-poke to the magazine was less than 2 seconds, no bait was placed in the active hole from the next session. The nose-poke and magazine entry time points, session duration, latency time from nose-poke to magazine approach, time spent in the magazine, and the number of nose-pokes and magazine entries were recorded using the Med-PC IV software, and the time resolution was 10 ms.

1. Fig. 1a: After 3 days of magazine training, the fixed ratio of 1 (FR1) was performed for 54 session days (Fig. 1a). During the FR1 tasks, the left hole served as the active nose-poke hole and the right hole as the inactive hole for the first 29 days (Fig. 1a, days 1-29). Starting from day 30, the roles of the holes reversed, with the left becoming inactive and the right becoming active for the next 17 days (Fig. 1a, days 30-46). This reversal pattern continued at set intervals: on day 47, the left hole became active again for 4 days (Fig. 1a, days 47-50), followed by a final reversal on day 51, with the right hole becoming active for the last 4 days (Fig. 1a, days 51-54).
2. Fig. 4b: After 3 days of magazine training, the FR1 task was performed for 20 session days (Fig. 4b). We treated mice trained in the FR1 task with C21 (3 mg/kg, intraperitoneally) 30 min before the nose-poke test on days 10 and 18. During the FR1 tasks, the left hole served as the active nose-poke hole and the right hole as the inactive hole for the first 9 days (Fig. 4b, days 1-9). Starting from day 10, the roles of the holes reversed, with the left becoming inactive and the right becoming active for the next 4 days (Fig. 4b, days 10-13). This reversal pattern continued at set intervals: on day 14, the left hole became active again for 4 days (Fig. 4b, days 14-17), followed by a final reversal on day 18, with the right hole becoming active for the last 3 days (Fig. 4b, days 18-20).
3. Fig. 5a: After 3 days of magazine training, the FR1 task was performed for 13 sessions (Fig. 5a). During the FR1 tasks, the left hole served as the active nose-poke hole and the right hole as the inactive hole for the first 9 days (Fig. 5a, days 1-9). Starting from day 10, the roles of the holes reversed, with the left becoming inactive and the right becoming active for the last 4 days (Fig. 5a, days 10-13).
4. Fig. 6c: After 3 days of magazine training, the FR1 task was performed for 13 sessions (Fig. 6c). We trained mice on the FR1 task for 7- and 10-days during nose-poke testing. The light was delivered using 2-ms pulse width, 10 Hz to ChR2 or eNpHR3.0, and the power measured at the tip of a 200-μm optical fiber was ∼3 mW. During the FR1 tasks, the left hole served as the active nose-poke hole and the right hole as the inactive hole for the first 9 days (Fig. 6c, days 1-9). Starting from day 10, the roles of the holes reversed, with the left becoming inactive and the right becoming active for the last 4 days (Fig. 6c, days 10-13).
5. Fig. 7c: After 3 days of magazine training, the FR1 task was performed for 13 session days (Fig. 7c). We treated mice trained in the FR1 task with C21 (3 mg/kg, intraperitoneally) 30 min before the nose-poke test on days 7 and 10. During the FR1 tasks, the left hole served as the active nose-poke hole and the right hole as the inactive hole for the first 9 days (Fig. 7c, days 1-9). Starting from day 10, the roles of the holes reversed, with the left becoming inactive and the right becoming active for the last 4 days (Fig. 7c, days 10-13).

### DeepLabCut

Markerless pose estimation was performed with the DeepLabCut toolbox (v.2.2.0.6)^56^. Six key points (canula, left ear, right ear, left hip, right hip, and tail base) of each animal were localized on each frame. In total, 3680 labeled frames were selected across 266 video recordings and used to train a deep learning model based on ResNet-50 with default parameters. Randomly assigned 95% of the data were used for training and the rest for testing. The network was trained for 1,030,000 iterations until cross-entropy loss plateaued at 6.84×10^-3^. We validated with 5 number of shuffles, and found the test error was: 4.86 pixels, train: 2.48 pixels (image size was 296 by 322). We then used a *p*- cutoff of 0.95 to condition the X and Y coordinates for future analysis. This network was then used to analyze videos from similar experimental settings.

### Session time-based learning phase division

To investigate how the interaction between GPe astrocytes and Proto^GPe→STN^ neurons changes throughout learning (Fig. 2d), while accounting for individual differences in learning speed, we divided each block into three distinct phases based on session time (Fig. 2e): “pre-learning” phase (L1/R1 in Fig. 2e), “learning” phase (L2/R2 in Fig. 2e), and “post-learning” phase (L3/R3 in Fig. 2e).

For each block, we modeled the session time for each subject across *L* days by fitting the function *f*, an exponential decay function with a step offset. The function *f* is defined as:

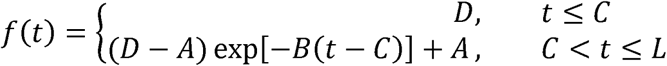

Until day *C*, the session time remains approximately constant at *J*. After day *C*, the session time begins to decay exponentially and eventually converges to *A*. We define the period from day 1 to day *C* as the pre-learning phase.

To determine the post-learning phase, where session times stabilize and stop decreasing, we applied a plateau-detection method^87, 88^. For the decaying portion of the fitted function (*C* ≤ *t* ≤ *L*), we drew a line *l*, connecting the initial point at *t*=*C* and the endpoint at *t*= *L*. We then defined the criterion *t*\* as the point where the derivative of the function *f* is closest to *s*, the slope of the line *l* drawn on *f*:

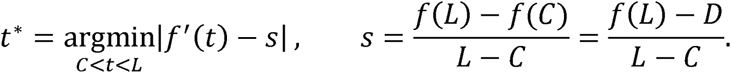

The learning phase is defined as the period from day *C*+ 1 to day *t*\*, and the post-learning phase is defined as the period from day *t*\* +1 to day *L*.

### Classification analysis

To investigate the context-related information in Ca^2+^ activity of GPe astrocyte or Proto^GPe→STN^ neurons around the magazine, we employed a support vector machine (SVM) using the “fitcsvm” function in MATLAB. We utilized a linear kernel for its minimal risk of overfitting and demonstrated capability in analyzing context-related cellular activity^89–92^.

The “fitcsvm” function was used with default parameters, including the regularization parameter, box constraint (set to 1), and the kernel scale parameter (set automatically for automatic scaling). Ca^2+^ activity during 2-s periods immediately before and after the magazine events (magazine entry and magazine exit) was used as input. ΔF/F0 data from different mice (*n* = 5) were normalized to z-scores to standardize the signal across subjects and sessions, mitigating the effects of potential baseline fluorescence drift and inter-subject variability in signal amplitude, and subsequently pooled together. The number of labels was 2 (left or right). To balance the dataset and prevent overfitting, we performed random under-sampling, repeated 1000 times for each analysis. SVM performance measures, including accuracy, sensitivity (accuracy in the left context), specificity (accuracy in the right context), and area under the receiver operator characteristic curve (AUROC), were collected after each iteration. As accuracy increases, AUROC approaches 1 (AUROC of 0.5 = chance; AUROC of 1.0 = 100% accuracy).

To further assess the robustness of the contextual encoding quantified by the SVM classification, we performed a validation analysis using different Ca^2+^ input time windows around magazine entry (ME) and magazine exit (MX) events. We tested time windows of 0.5, 1, 1.5, and 2 seconds both before and after ME and MX events. For each time window, we repeated the SVM classification procedure as described above. Across all tested time windows, the SVM classifiers consistently achieved accuracy significantly above the chance level (Supplementary Table 1).

Classification accuracy was used to quantify the degree of contextual encoding. Linear SVM-based classification was performed in a multi-dimensional space, where each axis represented Ca^2+^ activity at specific time points around the action event. Accuracy exceeding chance indicated that Ca^2+^ traces are linearly separable and form distinct contextual clusters (left vs right). The classifier assigns traces to contexts based on their proximity to these clusters, with higher accuracy reflecting a greater difference in distance from the trace to the two clusters. In this Ca^2+^ activity space, proximity, defined by Euclidean distance, reflects the similarity between the corresponding Ca^2+^ traces. The distance to a cluster represents how well a trace aligns with the stereotypic Ca^2+^ activity pattern of a given context (i.e., average Ca^2+^ activity during that context). Thus, higher classification accuracy signifies a stronger alignment of Ca^2+^ activity to the stereotypic pattern of one context compared to the other, indicating a greater degree of contextual encoding.

### Definition of the trajectory regularity

To provide a descriptive measure of a subject’s alignment to the optimal policy, we introduced the ‘trajectory regularity,’ calculated for each session (Extended Data Fig. 4a). The trajectory regularity accounts for the previous observations that a subject following a more optimal policy will show more stereotyped trajectories^59, 60^, defined by a cycle of (1) magazine exit (MX), (2) nose-poke (NP), and (3) magazine entry (ME), without additional variance introduced by unnecessary actions such as wandering.

During one session *S*, suppose that the mouse performs *N* cycles (MX-NP-ME). Each cycle *T_i_* can be represented as a 2D matrix of time points and spatial coordinates, where

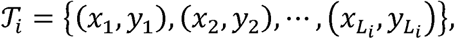

with *L_i_* being the number of time points in the cycle *T_i_* .

Since these cycles may vary in length, we used dynamic time warping (DTW)^93^ to align two cycles, *T_i_* and *T_j_*. Then, we calculated the normalized DTW distance by calculating the pairwise DTW distance between *T_i_* and *T_j_* and dividing it by the length of the aligned cycle^94^. Since cycle lengths varied, we normalized the DTW distance by the length of the aligned cycle to ensure comparability across cycle pairs, thus reflecting shape dissimilarity independent of cycle duration. We repeated this procedure for every possible pair of cycles within session *S*, producing a set of distances 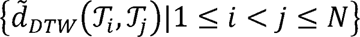.

The trajectory regularity for session *S* is then defined as the negative of the median of these pairwise DTW distances. A higher trajectory regularity value indicates more stereotyped and consistent trajectories across the session, whereas a lower trajectory regularity reflects more variable and less consistent behavior.

### Factor analysis to quantify behavioral optimality

To quantify behavioral optimality in the FR1 task, we conducted a factor analysis, rather than Principal Component Analysis (PCA), on three key behavioral metrics: session time (reflecting task efficiency), discrimination ratio (reflecting action selection accuracy), and trajectory regularity (reflecting motor consistency), established indicators of performance in operant conditioning. We chose factor analysis because our goal was to identify a latent variable representing ’behavioral optimality’ that explains the shared variance across these metrics, rather than simply reducing dimensionality, which is the primary aim of PCA. Furthermore, as our study involves longitudinal data with repeated measures over time, factor analysis offers advantages over PCA in mitigating the influence of time-dependent trends and noise, as recently demonstrated for temporal data analysis^95^.

Prior to factor analysis, each metric was z-score normalized for comparability. We then performed factor analysis to extract a single latent factor representing the shared variation across these metrics. This factor (Extended Data Fig. 4b) was interpreted as a composite measure of behavioral optimality, capturing the common variance reflecting efficient, accurate, and consistent task performance. Higher factor scores indicate greater behavior optimality.

### Validation of behavioral optimality factor using Leave-One-Out factor analysis

To further validate the robustness of our behavioral optimality factor, we implemented a leave-one-out factor analysis approach. For each of the three behavioral metrics (session time, discrimination ratio, and trajectory regularity), we iteratively “hid” one metric and performed factor analysis using only the remaining two metrics. Specifically, in each iteration: (1) one metric was designated as the ’hidden’ measure, (2) factor analysis was conducted on the other two metrics to extract a single latent factor, and (3) the correlation between the ’hidden’ measure and the extracted latent factor scores was calculated. This process was repeated three times, with each of the three behavioral metrics serving as the ’hidden’ measure in turn. Significant positive correlations across all three iterations would indicate that each individual behavioral metric consistently aligns with a latent factor derived from the other two, supporting the validity and robustness of the overall behavioral optimality factor.

To address the possibility that the robustness of the behavioral optimality factor might be influenced by general trends over time-related to experimental experience (i.e., “Day After Reversal”), we refined our leave-one-out factor analysis by first accounting for the “Day After Reversal” trend from each behavioral metric. For each metric (session time, discrimination ratio, and trajectory regularity), we performed a linear mixed model to predict its value based on “Day After Reversal”, while considering the subject as a random factor. We then calculated the residuals from these regressions. These residuals represent the variance in each metric that is not explained by the linear trend of “Day After Reversal,” effectively removing the time-dependent component.

Subsequently, we applied the leave-one-out factor analysis to these residualized behavioral metrics. Given the longitudinal nature of our study design, with repeated measures taken on the same subjects over 54 days, a standard Pearson correlation would be inappropriate as it assumes independence of observations, which is violated in our data. Therefore, the statistical significance of the correlations between the ’hidden’ residualized metrics and the latent factor scores was assessed using repeated-measures correlations^61^, a statistical technique specifically designed for longitudinal data that appropriately accounts for inter-subject variability and within-subject dependencies across repeated measurements.

### Statistical analyses

All data are represented as mean ± SEM using Prism 9.0 (GraphPad Software, San Diego, CA) and MATLAB R2023a (The Mathworks, Inc., Natick, MA). All data were analyzed by one-sample Wilcoxon signed-rank test, two-sample paired Wilcoxon signed-rank test, Mann-Whitney U test, mixed-effects logistic regression, mixed-effects linear regression, and cluster-based permutation test using MATLAB R2023a (The Mathworks, Inc., Natick, MA) and R (Ver. 4.1.2, R Core Team, R Foundation for Statistical Computing, Vienna, Austria). The statistical significance level was set at *P* < 0.05. The number of permutations for all cluster-based permutation tests was set at N = 10^4^. Detailed statistical tests and data with exact p values are listed in Supplementary Tables 1-3.

## Data availability

Upon publication, Source data will be provided in this paper. All raw data are available from the authors upon reasonable request.

## Code availability

Upon publication, all codes and preprocessed data used in this manuscript will be available in the public repository.

## Acknowledgments

We thank all the laboratory members for their helpful discussion and comments. We especially thank Dr. Ju Young Lee and Dr. Minryung Song for their insightful comments. Figures were created with BioRender.com. This research was supported by the Samuel C. Johnson for Genomics of Addiction Program at the Mayo Clinic, the Ulm Foundation, and the National Institute of Health (AA029258, AG072898, AA027773, MH137204). This study was supported by the National Research Foundation of Korea (NRF) grant funded by the Korean government (MSIT) (NRF-2019M3E5D2A01066267, Development of metacognitive AI for rapid learning), Electronics and Telecommunications Research Institute (ETRI) grant funded by the Korean government (22ZS1100, Core Technology Research for Self-Improving Integrated Artificial Intelligence System), and the Institute of Information & Communications Technology Planning & Evaluation (IITP) grant funded by the Korea government (MSIT) (No. 2019- 0-00075, Artificial Intelligence Graduate School Program (KAIST) and No. RS-2023- 00233251, System3 reinforcement learning with high-level brain functions). This research was supported by the Soonchunhyang University (2025-007).

## Author Contributions

M.A.Y., S.W.K. contributed equally to this work as the first author. M.A.Y., S.W.K., S.W.L., and D.S.C. designed the study. S.W.K. performed all the behavioral experiments. A.B. and S.K. performed all the *in vivo* electrophysiology experiments. M.A.Y. defined necessary concepts, analyzed all the data, and conducted all the statistical analyses. M.A.Y., S.W.K., S.W.L., and D.S.C. wrote and edited the manuscript. All authors revised and approved the paper.

## References

1. Uddin, L.Q. Cognitive and behavioural flexibility: neural mechanisms and clinical considerations. Nat Rev Neurosci 22, 167–179 (2021).

2. Uddin, L.Q. Brain Mechanisms Supporting Flexible Cognition and Behavior in Adolescents With Autism Spectrum Disorder. Biol Psychiatry 89, 172–183 (2021).

3. Duan, C.A., Erlich, J.C. & Brody, C.D. Requirement of Prefrontal and Midbrain Regions for Rapid Executive Control of Behavior in the Rat. Neuron 86, 1491–1503 (2015).

4. Reinert, S., Hubener, M., Bonhoeffer, T. & Goltstein, P.M. Mouse prefrontal cortex represents learned rules for categorization. Nature 593, 411–417 (2021).

5. Spellman, T., Svei, M., Kaminsky, J., Manzano-Nieves, G. & Liston, C. Prefrontal deep projection neurons enable cognitive flexibility via persistent feedback monitoring. Cell 184, 2750–2766 e2717 (2021).

6. Gangal, H., et al. Drug reinforcement impairs cognitive flexibility by inhibiting striatal cholinergic neurons. Nat Commun 14, 3886 (2023).

7. Li, H., et al. Silencing dentate newborn neurons alters excitatory/inhibitory balance and impairs behavioral inhibition and flexibility. Sci Adv 10, eadk4741 (2024).

8. Graybiel, A.M., Aosaki, T., Flaherty, A.W. & Kimura, M. The basal ganglia and adaptive motor control. Science 265, 1826–1831 (1994).

9. Okada, K., et al. Enhanced flexibility of place discrimination learning by targeting striatal cholinergic interneurons. Nat Commun 5, 3778 (2014).

10. Lilascharoen, V., et al. Divergent pallidal pathways underlying distinct Parkinsonian behavioral deficits. Nat Neurosci 24, 504–515 (2021).

11. Woolley, S.C., Rajan, R., Joshua, M. & Doupe, A.J. Emergence of context-dependent variability across a basal ganglia network. Neuron 82, 208–223 (2014).

12. Stephenson-Jones, M., et al. A basal ganglia circuit for evaluating action outcomes. Nature 539, 289–293 (2016).

13. Amita, H., Kim, H.F., Smith, M.K., Gopal, A. & Hikosaka, O. Neuronal connections of direct and indirect pathways for stable value memory in caudal basal ganglia. Eur J Neurosci 49, 712–725 (2019).

14. Larry, N., Zur, G. & Joshua, M. Organization of reward and movement signals in the basal ganglia and cerebellum. Nat Commun 15, 2119 (2024).

15. Kunimatsu, J., Suzuki, T.W., Ohmae, S. & Tanaka, M. Different contributions of preparatory activity in the basal ganglia and cerebellum for self-timing. Elife 7 (2018).

16. Khilkevich, A., et al. Brain-wide dynamics linking sensation to action during decision-making. Nature 634, 890–900 (2024).

17. Yin, H.H. & Knowlton, B.J. The role of the basal ganglia in habit formation. Nat Rev Neurosci 7, 464–476 (2006).

18. van Elzelingen, W., et al. Striatal dopamine signals are region specific and temporally stable across action-sequence habit formation. Curr Biol 32, 1163–1174 e1166 (2022).

19. Seiler, J.L., et al. Dopamine signaling in the dorsomedial striatum promotes compulsive behavior. Curr Biol 32, 1175–1188 e1175 (2022).

20. Huang, Z., et al. Dynamic responses of striatal cholinergic interneurons control behavioral flexibility. Sci Adv 10, eadn2446 (2024).

21. Xiao, L., Bornmann, C., Hatstatt-Burkle, L. & Scheiffele, P. Regulation of striatal cells and goal-directed behavior by cerebellar outputs. Nat Commun 9, 3133 (2018).

22. Shen, W., Flajolet, M., Greengard, P. & Surmeier, D.J. Dichotomous dopaminergic control of striatal synaptic plasticity. Science 321, 848–851 (2008).

23. Ragozzino, M.E., Ragozzino, K.E., Mizumori, S.J. & Kesner, R.P. Role of the dorsomedial striatum in behavioral flexibility for response and visual cue discrimination learning. Behav Neurosci 116, 105–115 (2002).

24. Haluk, D.M. & Floresco, S.B. Ventral striatal dopamine modulation of different forms of behavioral flexibility. Neuropsychopharmacology 34, 2041–2052 (2009).

25. Gittis, A.H., et al. New roles for the external globus pallidus in basal ganglia circuits and behavior. J Neurosci 34, 15178–15183 (2014).

26. Courtney, C.D. & Chan, C.S. Cell type-specific processing of non-motor signals in the external pallidum. Trends Neurosci 46, 336–337 (2023).

27. Albin, R.L., Young, A.B. & Penney, J.B. The functional anatomy of basal ganglia disorders. Trends Neurosci 12, 366–375 (1989).

28. Redgrave, P., et al. Goal-directed and habitual control in the basal ganglia: implications for Parkinson’s disease. Nat Rev Neurosci 11, 760–772 (2010).

29. Penney, J.B., Jr. & Young, A.B. Striatal inhomogeneities and basal ganglia function. Mov Disord 1, 3–15 (1986).

30. DeLong, M.R. Primate models of movement disorders of basal ganglia origin. Trends Neurosci 13, 281–285 (1990).

31. Mastrogiacomo, R., et al. Dysbindin-1A modulation of astrocytic dopamine and basal ganglia dependent behaviors relevant to schizophrenia. Mol Psychiatry 27, 4201–4217 (2022).

32. Johansson, Y. & Ketzef, M. Sensory processing in external globus pallidus neurons. Cell Rep 42, 111952 (2023).

33. Katabi, S., Adler, A., Deffains, M. & Bergman, H. Dichotomous activity and function of neurons with low- and high-frequency discharge in the external globus pallidus of non-human primates. Cell Rep 42, 111898 (2023).

34. Cui, Q., et al. Dissociable Roles of Pallidal Neuron Subtypes in Regulating Motor Patterns. J Neurosci 41, 4036–4059 (2021).

35. Nambu, A., Tokuno, H. & Takada, M. Functional significance of the cortico-subthalamo-pallidal ’hyperdirect’ pathway. Neurosci Res 43, 111–117 (2002).

36. Dunovan, K., Lynch, B., Molesworth, T. & Verstynen, T. Competing basal ganglia pathways determine the difference between stopping and deciding not to go. Elife 4, e08723 (2015).

37. Polyakova, Z., Chiken, S., Hatanaka, N. & Nambu, A. Cortical Control of Subthalamic Neuronal Activity through the Hyperdirect and Indirect Pathways in Monkeys. J Neurosci 40, 7451–7463 (2020).

38. Guillaumin, A., Serra, G.P., Georges, F. & Wallen-Mackenzie, A. Experimental investigation into the role of the subthalamic nucleus (STN) in motor control using optogenetics in mice. Brain Res 1755, 147226 (2021).

39. Aristieta, A., et al. A Disynaptic Circuit in the Globus Pallidus Controls Locomotion Inhibition. Curr Biol 31, 707–721 e707 (2021).

40. Ketzef, M. & Silberberg, G. Differential Synaptic Input to External Globus Pallidus Neuronal Subpopulations In Vivo. Neuron 109, 516–529 e514 (2021).

41. Hernandez, V.M., et al. Parvalbumin+ Neurons and Npas1+ Neurons Are Distinct Neuron Classes in the Mouse External Globus Pallidus. J Neurosci 35, 11830–11847 (2015).

42. Mallet, N., et al. Dichotomous organization of the external globus pallidus. Neuron 74, 1075–1086 (2012).

43. Crompe, B., et al. The globus pallidus orchestrates abnormal network dynamics in a model of Parkinsonism. Nat Commun 11, 1570 (2020).

44. Isett, B.R., et al. The indirect pathway of the basal ganglia promotes transient punishment but not motor suppression. Neuron 111, 2218–2231 e2214 (2023).

45. Cui, Q., et al. Blunted mGluR Activation Disinhibits Striatopallidal Transmission in Parkinsonian Mice. Cell Rep 17, 2431–2444 (2016).

46. Yu, X., et al. Context-Specific Striatal Astrocyte Molecular Responses Are Phenotypically Exploitable. Neuron 108, 1146–1162 e1110 (2020).

47. Lee, J.H., et al. Astrocytes phagocytose adult hippocampal synapses for circuit homeostasis. Nature 590, 612–617 (2021).

48. Kruyer, A., Dixon, D., Angelis, A., Amato, D. & Kalivas, P.W. Astrocytes in the ventral pallidum extinguish heroin seeking through GAT-3 upregulation and morphological plasticity at D1-MSN terminals. Mol Psychiatry 27, 855–864 (2022).

49. Soto, J.S., et al. Astrocyte-neuron subproteomes and obsessive-compulsive disorder mechanisms. Nature 616, 764–773 (2023).

50. Ollivier, M., et al. Crym-positive striatal astrocytes gate perseverative behaviour. Nature 627, 358–366 (2024).

51. Sheikhbahaei, S., et al. Astrocytes modulate brainstem respiratory rhythm-generating circuits and determine exercise capacity. Nat Commun 9, 370 (2018).

52. Lines, J., Martin, E.D., Kofuji, P., Aguilar, J. & Araque, A. Astrocytes modulate sensory-evoked neuronal network activity. Nat Commun 11, 3689 (2020).

53. Cho, W.H., et al. Hippocampal astrocytes modulate anxiety-like behavior. Nat Commun 13, 6536 (2022).

54. Kang, S., et al. Activation of Astrocytes in the Dorsomedial Striatum Facilitates Transition From Habitual to Goal-Directed Reward-Seeking Behavior. Biol Psychiatry 88, 797–808 (2020).

55. Kang, S., et al. Astrocyte activities in the external globus pallidus regulate action-selection strategies in reward-seeking behaviors. Sci Adv 9, eadh9239 (2023).

56. Mathis, A., et al. DeepLabCut: markerless pose estimation of user-defined body parts with deep learning. Nat Neurosci 21, 1281–1289 (2018).

57. Chereau, R., et al. Dynamic perceptual feature selectivity in primary somatosensory cortex upon reversal learning. Nat Commun 11, 3245 (2020).

58. Sun, J., Wang, S., Zhang, J. & Zong, C. Neural Encoding and Decoding With Distributed Sentence Representations. IEEE Trans Neural Netw Learn Syst 32, 589–603 (2021).

59. Dhawale, A.K., Wolff, S.B.E., Ko, R. & Olveczky, B.P. The basal ganglia control the detailed kinematics of learned motor skills. Nat Neurosci 24, 1256–1269 (2021).

60. Wolff, S.B.E., Ko, R. & Olveczky, B.P. Distinct roles for motor cortical and thalamic inputs to striatum during motor skill learning and execution. Sci Adv 8, eabk0231 (2022).

61. Bakdash, J.Z. & Marusich, L.R. Repeated Measures Correlation. Frontiers in Psychology 8 (2017).

62. Smart, A.D., et al. Engineering a light-activated caspase-3 for precise ablation of neurons in vivo. Proc Natl Acad Sci U S A 114, E8174–E8183 (2017).

63. Baker, M., et al. External globus pallidus input to the dorsal striatum regulates habitual seeking behavior in male mice. Nat Commun 14, 4085 (2023).

64. Courtney, C.D., Pamukcu, A. & Chan, C.S. Cell and circuit complexity of the external globus pallidus. Nat Neurosci 26, 1147–1159 (2023).

65. Serra, G.P., et al. A role for the subthalamic nucleus in aversive learning. Cell Rep 42, 113328 (2023).

66. Bergstrom, H.C., et al. Dorsolateral Striatum Engagement Interferes with Early Discrimination Learning. Cell Rep 23, 2264–2272 (2018).

67. Turner, K.M., Svegborn, A., Langguth, M., McKenzie, C. & Robbins, T.W. Opposing Roles of the Dorsolateral and Dorsomedial Striatum in the Acquisition of Skilled Action Sequencing in Rats. J Neurosci 42, 2039–2051 (2022).

68. Murphy-Royal, C., Ching, S. & Papouin, T. A conceptual framework for astrocyte function. Nat Neurosci 26, 1848–1856 (2023).

69. Oe, Y., et al. Distinct temporal integration of noradrenaline signaling by astrocytic second messengers during vigilance. Nat Commun 11, 471 (2020).

70. Corkrum, M., et al. Dopamine-Evoked Synaptic Regulation in the Nucleus Accumbens Requires Astrocyte Activity. Neuron 105, 1036–1047 e1035 (2020).

71. Vardjan, N., Parpura, V. & Zorec, R. Loose excitation-secretion coupling in astrocytes. Glia 64, 655–667 (2016).

72. Abdi, A., et al. Prototypic and arkypallidal neurons in the dopamine-intact external globus pallidus. J Neurosci 35, 6667–6688 (2015).

73. Dodson, P.D., et al. Distinct developmental origins manifest in the specialized encoding of movement by adult neurons of the external globus pallidus. Neuron 86, 501–513 (2015).

74. Saunders, A., et al. A direct GABAergic output from the basal ganglia to frontal cortex. Nature 521, 85–89 (2015).

75. Abecassis, Z.A., et al. Npas1(+)-Nkx2.1(+) Neurons Are an Integral Part of the Cortico-pallido-cortical Loop. J Neurosci 40, 743–768 (2020).

76. Ahrlund-Richter, S., et al. A whole-brain atlas of monosynaptic input targeting four different cell types in the medial prefrontal cortex of the mouse. Nat Neurosci 22, 657–668 (2019).

77. Foster, N.N., et al. The mouse cortico-basal ganglia-thalamic network. Nature 598, 188–194 (2021).

78. Poulin, J.F., et al. Mapping projections of molecularly defined dopamine neuron subtypes using intersectional genetic approaches. Nat Neurosci 21, 1260–1271 (2018).

79. Azcorra, M., et al. Unique functional responses differentially map onto genetic subtypes of dopamine neurons. Nat Neurosci 26, 1762–1774 (2023).

80. Assous, M., et al. Differential processing of thalamic information via distinct striatal interneuron circuits. Nat Commun 8, 15860 (2017).

81. Kincaid, A.E., Penney, J.B., Jr., Young, A.B. & Newman, S.W. The globus pallidus receives a projection from the parafascicular nucleus in the rat. Brain Res 553, 18–26 (1991).

82. Alcaraz, F., et al. Thalamocortical and corticothalamic pathways differentially contribute to goal-directed behaviors in the rat. Elife 7 (2018).

83. Karube, F., Takahashi, S., Kobayashi, K. & Fujiyama, F. Motor cortex can directly drive the globus pallidus neurons in a projection neuron type-dependent manner in the rat. Elife 8 (2019).

84. Xu, X., Song, L., Kringel, R. & Hanganu-Opatz, I.L. Developmental decrease of entorhinal-hippocampal communication in immune-challenged DISC1 knockdown mice. Nat Commun 12, 6810 (2021).

85. Sailer, L.L., Park, A.H., Galvez, A. & Ophir, A.G. Lateral septum DREADD activation alters male prairie vole prosocial and antisocial behaviors, not partner preferences. Commun Biol 5, 1299 (2022).

86. Pochinok, I., Stober, T.M., Triesch, J., Chini, M. & Hanganu-Opatz, I.L. A developmental increase of inhibition promotes the emergence of hippocampal ripples. Nat Commun 15, 738 (2024).

87. Hoare, S.R.J., Tewson, P.H., Quinn, A.M., Hughes, T.E. & Bridge, L.J. Analyzing kinetic signaling data for G-protein-coupled receptors. Sci Rep 10, 12263 (2020).

88. Luuk Loeff and Jacob, W.J.K.a.C.J.a.C.D. AutoStepfinder: A fast and automated step detection method for single-molecule analysis. Patterns 2, 100256 (2021).

89. Duarte, J.M., et al. Hippocampal contextualization of social rewards in mice. Nat Commun 15, 9493 (2024).

90. Ehret, B., et al. Population-level coding of avoidance learning in medial prefrontal cortex. Nat Neurosci 27, 1805–1815 (2024).

91. Elston, T.W. & Wallis, J.D. Context-dependent decision-making in the primate hippocampal–prefrontal circuit. Nature Neuroscience (2025).

92. Mishchanchuk, K., et al. Hidden state inference requires abstract contextual representations in the ventral hippocampus. Science 386, 926–932 (2024).

93. Zhang, Y., et al. Detailed mapping of behavior reveals the formation of prelimbic neural ensembles across operant learning. Neuron 110, 674–685.e676 (2022).

94. Müller, M. *Fundamentals of music processing: Audio, analysis, algorithms, applications* (Springer, 2015).

95. Francois, O. & Jay, F. Factor analysis of ancient population genomic samples. Nat Commun 11, 4661 (2020).

