## Extended Data Fig for "Pallidal prototypic neuron and astrocyte activities regulate flexible reward-seeking behaviors"

[1]Department of Molecular Pharmacology and Experimental Therapeutics, Mayo Clinic College of Medicine and Science, Rochester, Minnesota 55905, USA, [2]Department of Clinical Pharmacology, College of Medicine, Soonchunhyang University, 31151, Cheonan-si; [3]Department of Bio and Brain Engineering; [4]Program of Brain and Cognitive Engineering, Korea Advanced Institute of Science and Technology (KAIST), 34141, Daejeon, Republic of Korea, [5]Department of Pharmacology and Toxicology, Medical College of Georgia, Augusta University, Augusta, Georgia 30912, USA, [6]Department of Brain & Cognitive Sciences; [7]Kim Jaechul Graduate School of AI, Korea Advanced Institute of Science and Technology (KAIST), 34141, Daejeon, Republic of Korea, [8]Department of Psychiatry and Psychology; [9]Neuroscience Program, Mayo Clinic College of Medicine and Science, Rochester, Minnesota 55905, USA.

[†]These authors contributed equally to this work as the first author.

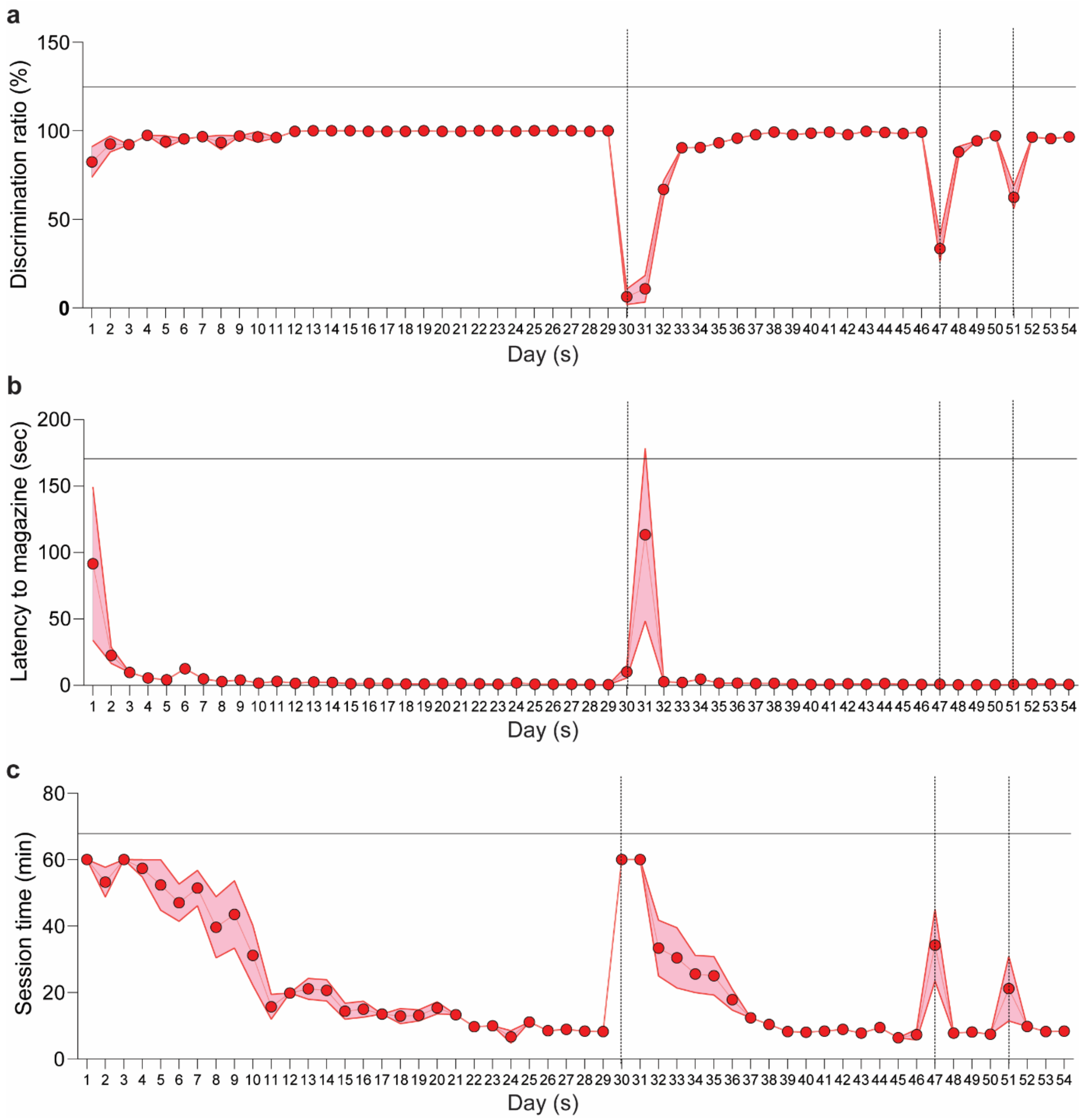


**Extended Data Fig. 1. Behavioral metrics during FR1 task training.** Vertical dashed lines indicate reversal days. **a,** Discrimination ratio (%) between active and inactive nose-pokes across training days. **b,** Latency (seconds) from active nose-poke to magazine entry across training days. **c,** Session duration (minutes) across training days. Shaded areas represent SEM. Data consist of a total of 261 sessions, collected across four blocks from 5 mice: *n* = 140 (1$^{st}$ block), 82 (2$^{nd}$ block), 19 (3$^{rd}$ block), and 20 (4$^{th}$ block).

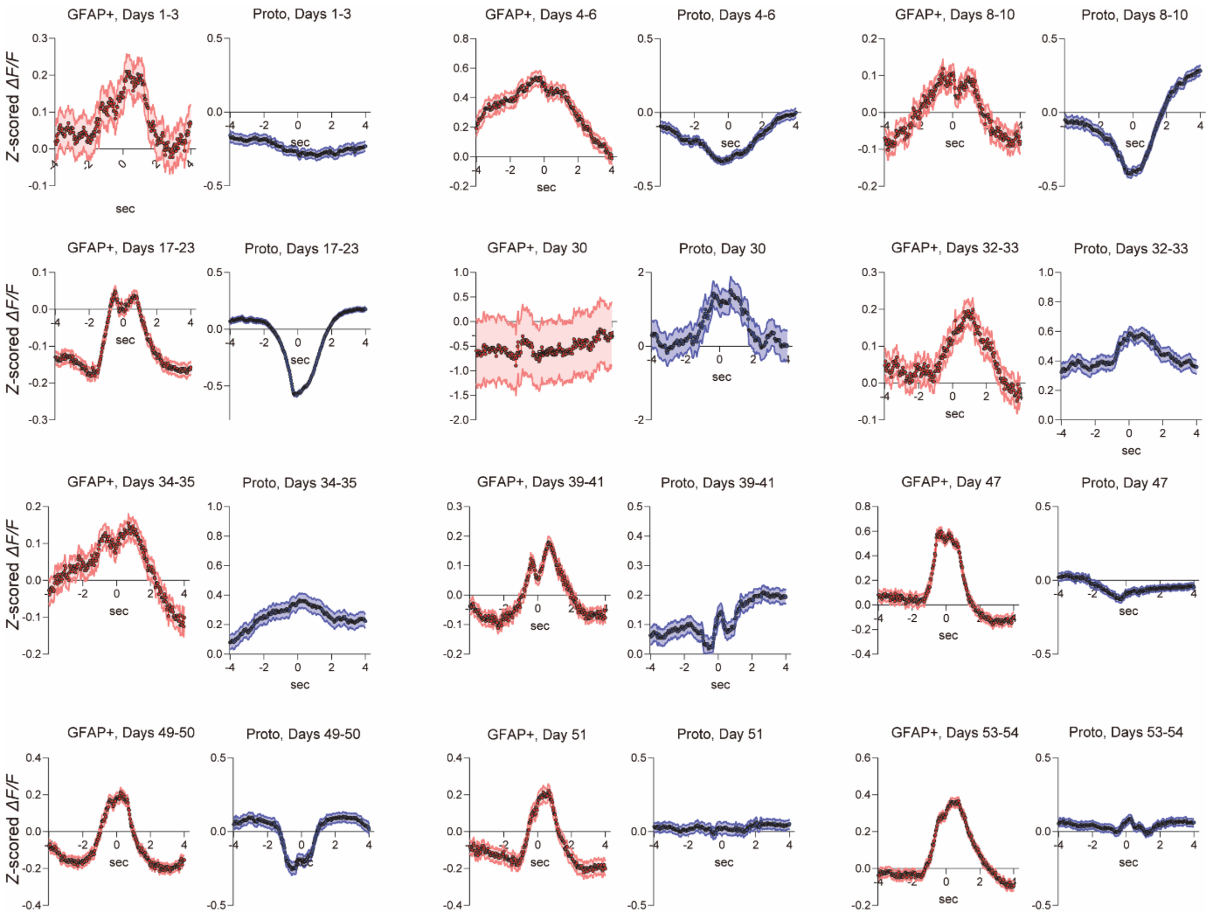


**Extended Data Fig. 2. Longitudinal calcium activity dynamics of GPe astrocytes and Proto$^{GPe\rightarrow STN}$ neurons.** Longitudinal calcium activity dynamics of GPe astrocytes (GFAP+, red) and Proto$^{GPe\rightarrow STN}$ neurons (Proto, blue) around active nose-pokes, aligned to the active nose-poke (0 s). Shaded areas represent SEM. Each panel corresponds to a specific task period, comparing GPe astrocyte and Proto$^{GPe\rightarrow STN}$ neuron activity. *n* = 5 mice.

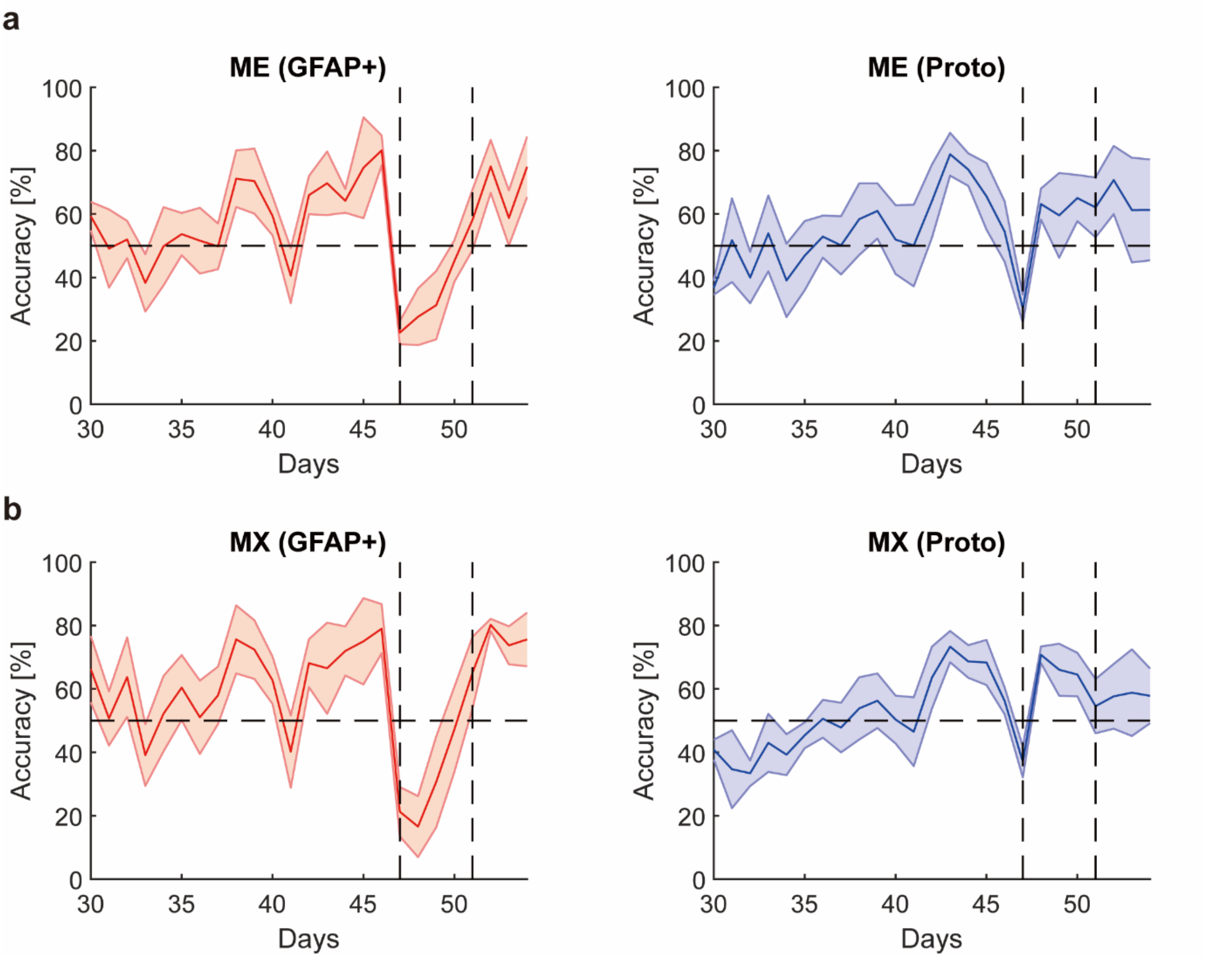


**Extended Data Fig. 3. Changes in classification accuracy across days following the 2nd and 3rd reversals.** Vertical dashed lines mark reversal days. **a,** Classification accuracy for models trained on $Ca^{2+}$ traces of GPe astrocytes (GFAP+, left) and $Proto^{GPe \rightarrow STN}$ neurons (Proto, right) around ME. **b,** Classification accuracy for models trained on $Ca^{2+}$ traces of GPe astrocytes (GFAP+, left) and $Proto^{GPe \rightarrow STN}$ neurons (Proto, right) around MX. Shaded areas represent SEM. Data consist of a total of 121 sessions, collected across three blocks from 5 mice; *n* = 82 (2nd block), 19 (3rd block), and 20 (4th block).

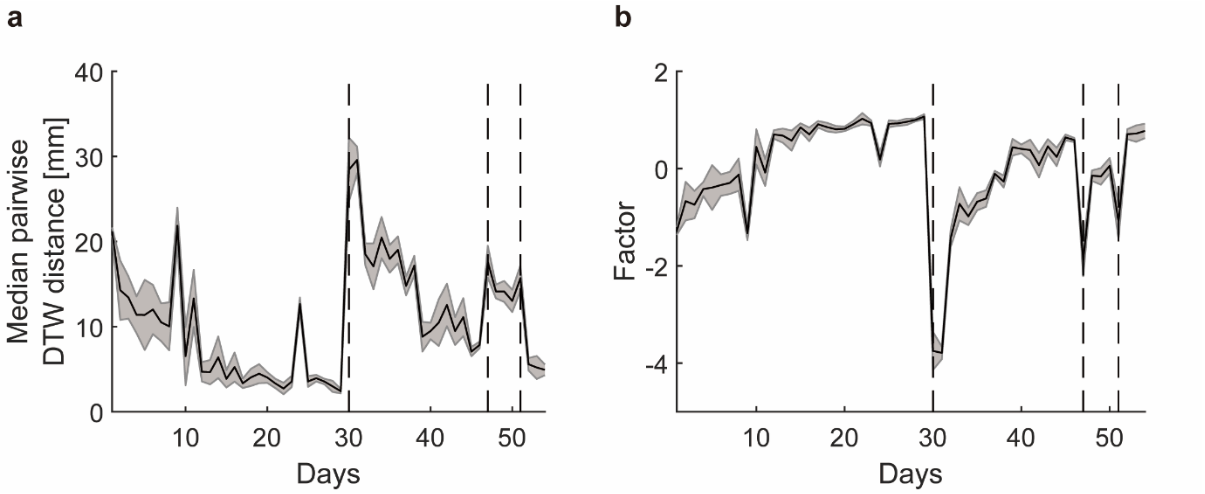


**Extended Data Fig. 4. Behavioral metrics reflecting behavioral optimality during FR1 task training.** Vertical dashed lines mark reversal days. **a,** Median pairwise DTW distance (mm) between reward acquisition cycles (active nose-poke, magazine entry/exit, and return to the active nose-poke hole) across training days. Smaller distances indicate reduced variation and higher trajectory regularity. **b,** Behavioral optimality factor derived via factor analysis of session time (task efficiency), discrimination ratio (accuracy), and trajectory regularity (consistency). Higher factor scores indicate more optimal behavior. Shaded areas represent SEM. Data consist of a total of 259 sessions, collected acorss four blocks from 5 mice; *n* = 138 ($1^{st}$ block), 82 ($2^{nd}$ block), 19 ($3^{rd}$ block), and 20 ($4^{th}$ block).

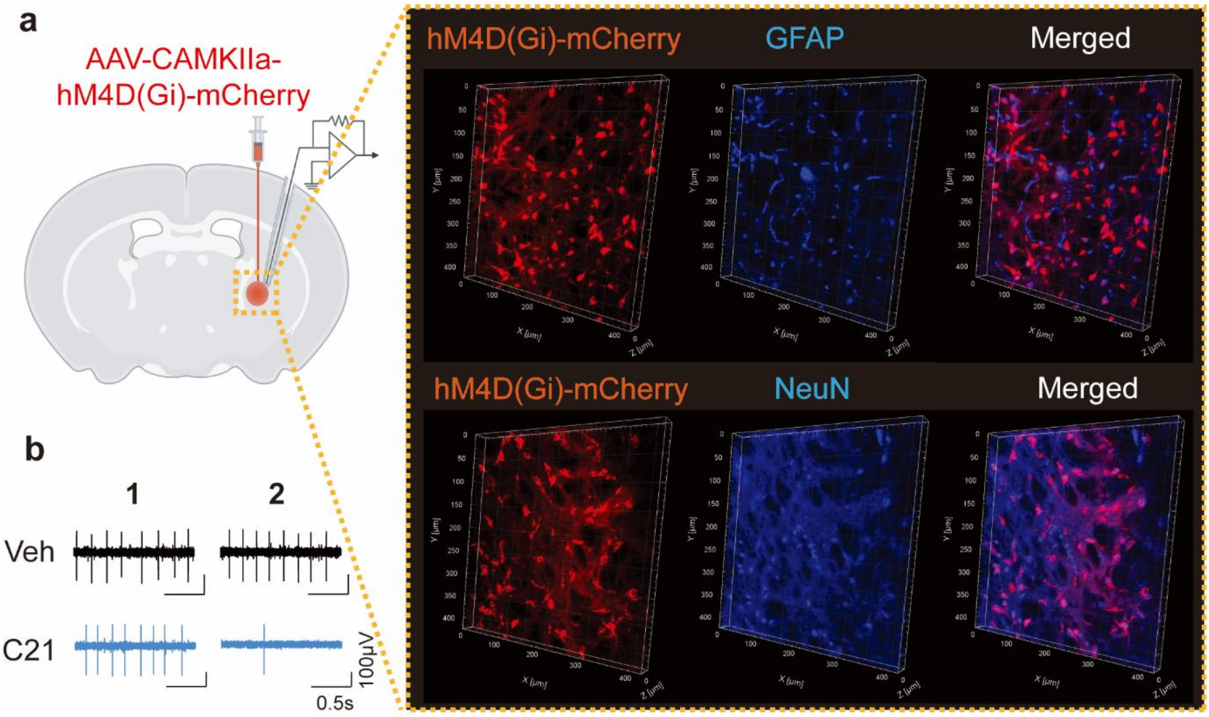


**Extended Data Fig. 5. Chemogenetic inhibition of GPe neurons via hM4D(Gi)-mCherry suppresses neuronal firing. a,** Representative images of hM4Di-mCherry expression in GPe neurons, colocalized with GFAP and NeuN. **b,** Representative electrophysiological traces demonstrating reduced neuronal firing with C21 (inhibitory DREADD ligand) compared to vehicle (Veh). $N_{Cell}$ = 21–23 per group (6 mice).

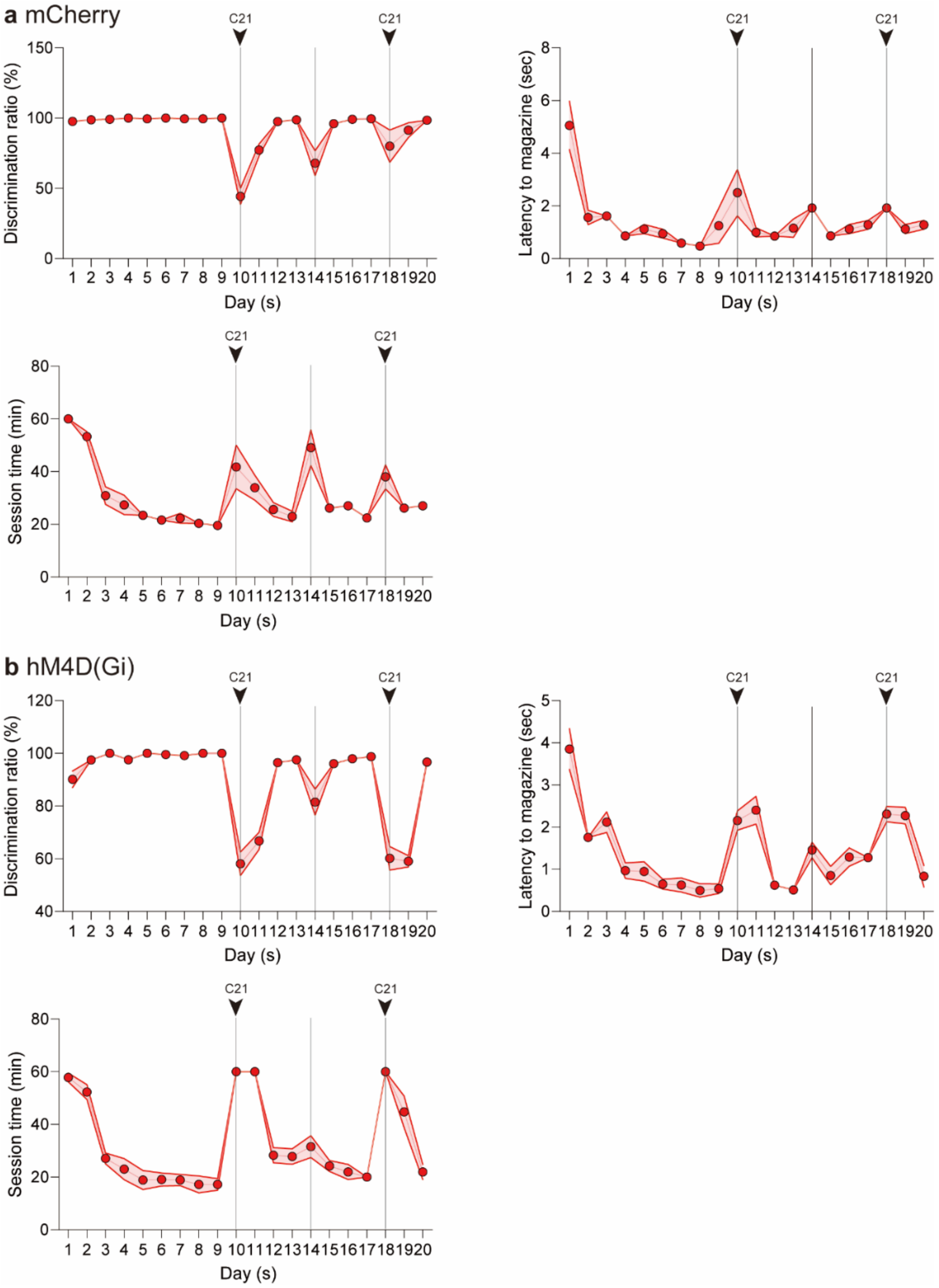


**Extended Data Fig. 6. Behavioral metrics during FR1 task training with chemogenetic inactivation of GPe neurons.** Vertical dashed lines mark reversal days, and arrows indicate C21 injection days (Days 9 and 15). Shaded areas represent SEM. Top left for discrimination ratio (%), Top right for latency (sec) from active nose-poke to magazine entry, Bottom for session time (min) across training days. **a,** Control group (mCherry). **b,** Inhibition group [hM4D(Gi)]. Data are represented as mean ± SEM; Data consist of behavioral performance of 5 mice per group, with identical session counts per block for both groups: *n* = 45 (1$^{st}$ block), 20 (2$^{nd}$ block), 20 (3$^{rd}$ block), and 15 (4$^{th}$ block).

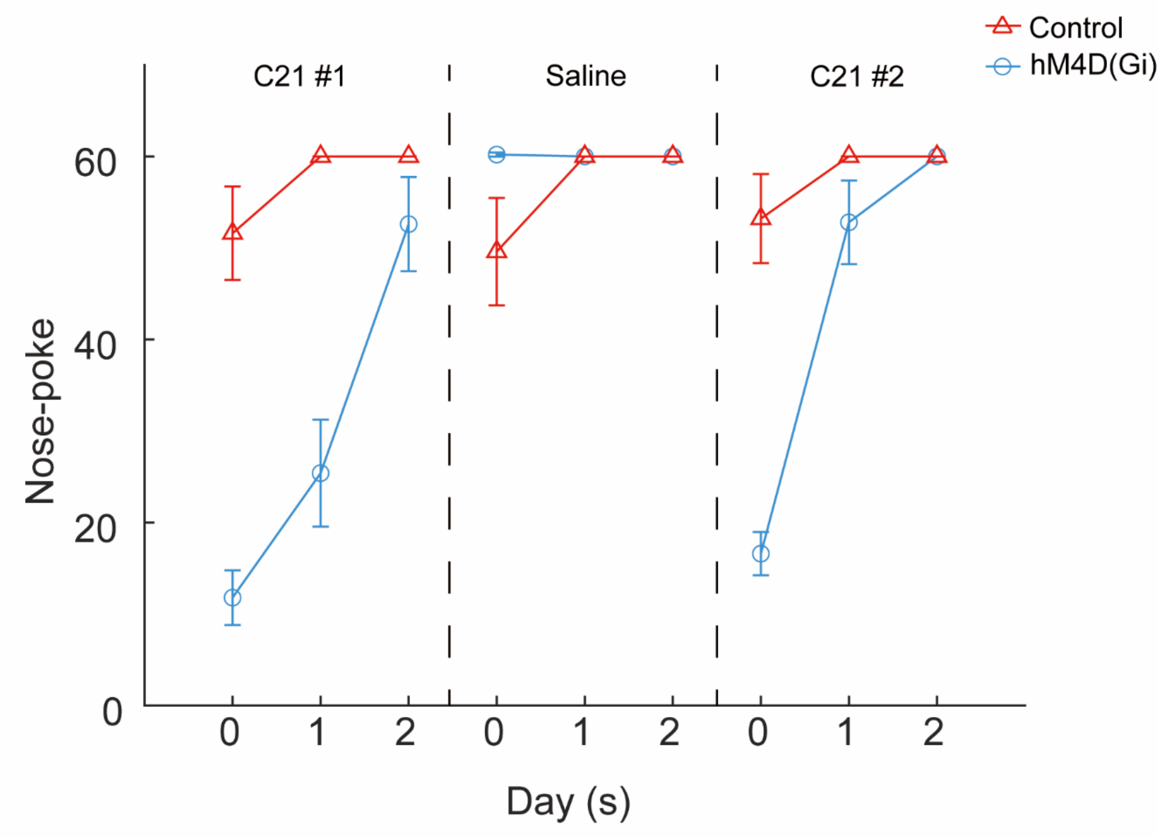


**Extended Data Fig. 7. Nose-poke behaviors after reversals with chemogenetic inactivation of GPe neurons.** Day 0 indicates the first day after reversal. Data are represented as mean ± SEM; Data consist of behavioral performance of 5 mice per group, with identical session counts per block for both groups: $n$ = 15 ($2^{nd}$ block), 15 ($3^{rd}$ block), and 15 ($4^{th}$ block). See Supplementary Table 1 for full statistical information.

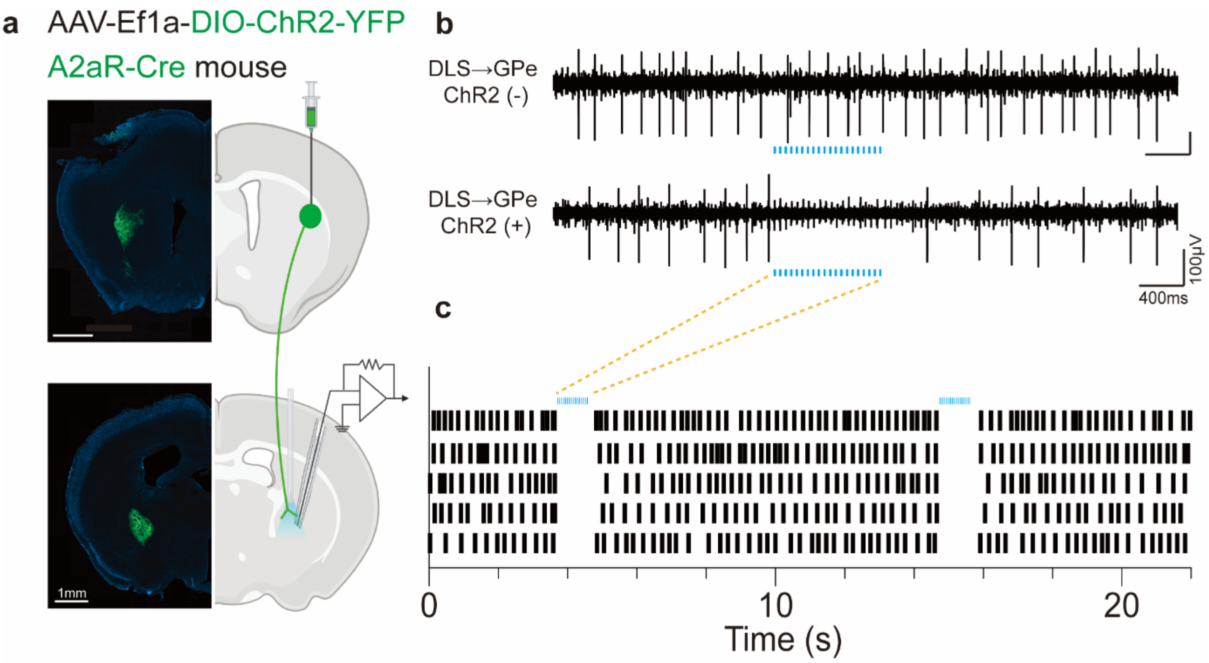


**Extended Data Fig. 8. Optogenetic stimulation of DLS neurons in the DLS→GPe circuit.** **a,** Circuit-dependent expression of ChR2-eYFP in A2aR-Cre mice, showing targeted labeling of the DLS→GPe pathway. **b,** Representative electrophysiological traces showing neuronal responses in the GPe with and without ChR2 activation. **c,** Raster plots illustrating neuronal responses to optogenetic excitation of synaptic terminals of A2A-positive DLS neurons in the GPe.

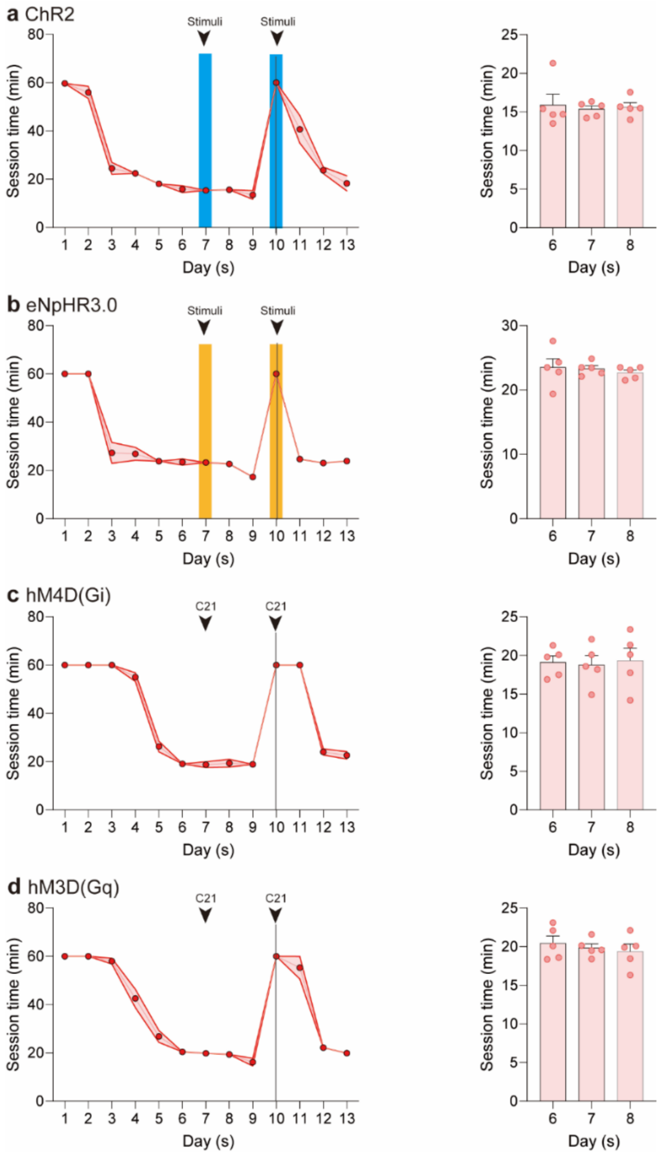


**Extended Data Fig. 9. Changes in session time during FR1 task training with optogenetic and chemogenetic manipulations.** Vertical dashed line marks the reversal day, and shaded areas represent SEM. Bar graphs on the right display session durations at critical intervention day (Day 7) compared to days before (Day 6) and after (Day 8) the intervention day. **a-b,** Session time changes with optogenetic interventions in STR-GPe pathways. **a,** Optogenetic activation (ChR2). **b,** Optogenetic inhibition (eNpHR3.0). **c-d,** Session time changes with chemogenetic manipulations in STN-GPe pathways. **c,** Chemogenetic inhibition [hM4D(Gi)]. **d,** Chemogenetic activation [hM3D(Gq)]. Data are represented as mean ± SEM; For optogenetic manipulation (**a-b**), data consist of behavioral performance of 5 mice per group, with identical session counts per block for both groups: $n$ = 45 ($1^{st}$ block), and 20 ($2^{nd}$ block). For chemogenetic manipulation (**c-d**), data consist of behavioral performance of 5 mice per group, with identical session counts per block for both groups: $n$ = 45 ($1^{st}$ block), and 20 ($2^{nd}$ block). See Supplementary Table 1 for full statistical information.
