## Supplementary Table for "Pallidal prototypic neuron and astrocyte activities regulate flexible reward-seeking behaviors"

**Supplementary Table 1. Summary of statistical analysis**

| Figure | | Statistical Tests | Comparison | Value & Results | *P* value |
|---|---|---|---|---|---|
| **Fig. 1** | **d** | Two-way ANOVA | Interaction | F (53, 432) = 58.250 | *p<0.0001* |
| | | | Time | F (53, 432) = 5.320 | *p<0.0001* |
| | | | Group | F (1, 432) = 223.2 | *p<0.0001* |
| **Fig. 2** | **d** | Two-way ANOVA | Interaction | F (240, 1928) = 0.430 | *p<0.0001* |
| | | | Raw Factor | F (240, 1928) = 0.8865 | *p=0.8849* |
| | | | Column Factor | F (1, 1928) = 7169 | *p<0.0001* |
| | | Two-way ANOVA | Interaction | F (240, 1928) = 34.75 | *p<0.0001* |
| | | | Raw Factor | F (240, 1928) = 12.84 | *p<0.0001* |
| | | | Column Factor | F (1, 1928) = 985.8 | *p<0.0001* |
| | | Two-way ANOVA | Interaction | F (240, 1928) = 47.38 | *p<0.0001* |
| | | | Raw Factor | F (240, 1928) = 39.36 | *p<0.0001* |
| | | | Column Factor | F (1, 1928) = 396.6 | *p<0.0001* |
| | | Two-way ANOVA | Interaction | F (240, 1928) = 66.60 | *p<0.0001* |
| | | | Raw Factor | F (240, 1928) = 19.86 | *p<0.0001* |
| | | | Column Factor | F (1, 1928) = 1054 | *p<0.0001* |
| | | Two-way ANOVA | Interaction | F (240, 1928) = 0.9402 | *p=0.7280* |
| | | | Raw Factor | F (240, 1928) = 0.6108 | *p>0.9999* |
| | | | Column Factor | F (1, 1928) = 594.1 | *p<0.0001* |
| | | Two-way ANOVA | Interaction | F (240, 1928) = 10.55 | *p<0.0001* |
| | | | Raw Factor | F (240, 1928) = 15.41 | *p<0.0001* |
| | | | Column Factor | F (1, 1928) = 3940 | *p<0.0001* |
| | | Two-way ANOVA | Interaction | F (240, 1928) = 9.713 | *p<0.0001* |
| | | | Raw Factor | F (240, 1928) = 16.92 | *p<0.0001* |
| | | | Column Factor | F (1, 1928) = 205.8 | *p<0.0001* |
| Figure 2f. We analyzed the change in average $Ca^{2+}$ activity difference at different stages using a linear mixed-effects model, with phase as the fixed factor and subject as a random effect. Coefficient significance was assessed after adjusting degrees of freedom using the Kenward-Roger method. Estimated marginal means were compared after Satterthwaite-adjusted degrees of freedom. Estimates are reported as mean ± SEM. | | | | | |

| | | | | | | |
|---|---|---|---|---|---|---|
| **Fig. 2** | **f** | Increasing trend (L1-L3) | 140 sessions from 5 mice | Repeated-measures correlation | r(134) = 0.2802, p = $9.5377\times10^{-4}$ | |
| | | Increase after reversal (L3-R1) | 86 sessions from 5 mice | Linear mixed-effects model | Phase: F(1,82.12) = 42.503, | *p*<0.001 |
| | | | | Main effect test for comparing estimated marginal means | L3: -0.127 ± 0.051<br>R1: 0.720 ± 0.121<br>R1-L3: 0.848 ± 0.129, t(82.129) = 6.571, p < 0.001 | |
| | | Decreasing trend (R1-R3) | 79 sessions from 5 [illegible] | Repeated-measures correlation | r(73) = -0.4977, p = $5.5411\times10^{-6}$ | |
| Figure 3c. Estimates are reported as mean ± bootstrap standard error, as they are based on random under-sampling. | | | | | | |
| **Fig. 3** | **c Top (ME)** | Accuracy across blocks (Overall) | 1000 random seeds | One-sample bootstrap hypothesis test against chance level | [GFAP+]: 71.46 ± 1.12%<br>[Proto]: 77.80 ± 1.03% | [GFAP+]: *p*<0.001<br>[Proto]: *p*<0.001 |
| | | | | Paired two-sample bootstrap | [GFAP+]-[Proto]: -6.34 ± 1.50% | *p*<0.001 |
| | | Accuracy in the left context (LNP) | 1000 random seeds | One-sample bootstrap hypothesis test against chance level | [GFAP+]: 64.95 ± 2.34%<br>[Proto]: 84.44 ± 1.26% | [GFAP+]: *p*<0.001<br>[Proto]: *p*<0.001 |
| | | | | Paired two-sample bootstrap | [GFAP+]-[Proto]: -19.50 ± 2.70% | *p*<0.001 |
| | | Accuracy in the right context (RNP) | 1000 random seeds | One-sample bootstrap hypothesis test against chance level | [GFAP+]: 77.97 ± 1.83%<br>[Proto]: 71.16 ± 1.64% | [GFAP+]: *p*<0.001<br>[Proto]: *p*<0.001 |
| | | | | Paired two-sample bootstrap hypothesis test | [GFAP+]-[Proto]: 6.81 ± 2.42% | *p*= 0.004 |
| | | AUROC | 1000 random seeds | One-sample bootstrap hypothesis test against chance [illegible] | [GFAP+]: 78.88 ± 1.02%<br>[Proto]: 85.32 ± 0.83% | [GFAP+]: *p*<0.001<br>[Proto]: *p*<0.001 |
| | | | | Paired two-sample bootstrap | [GFAP+]-[Proto]: -6.44 ± 1.29% | *p*<0.001 |

| | | | | | | |
|---|---|---|---|---|---|---|
| **Fig. 3** | **c Bottom (ME)** | Accuracy across blocks (Overall) | 1000 random seeds | One-sample bootstrap hypothesis test against chance level | [GFAP+]: 69.64 ± 1.13%<br>[Proto]: 74.89 ± 1.00% | [GFAP+]: $p<0.001$<br>[Proto]: $p<0.001$ |
| | | | | Paired two-sample bootstrap | [GFAP+]-[Proto]: -5.25 ± 1.48% | $p=0.004$ |
| | | Accuracy in the left context (LNP) | 1000 random seeds | One-sample bootstrap hypothesis test against chance level | [GFAP+]: 61.25 ± 1.99%<br>[Proto]: 80.75 ± 1.22% | [GFAP+]: $p<0.001$<br>[Proto]: $p<0.001$ |
| | | | | Paired two-sample bootstrap | [GFAP+]-[Proto]: -19.50 ± 2.38% | $p<0.0001$ |
| | | Accuracy in the right context (RNP) | 1000 random seeds | One-sample bootstrap hypothesis test against chance [illegible] | [GFAP+]: 78.03 ± 1.54%<br>[Proto]: 69.04 ± 1.60%, | [GFAP+]: $p<0.001$<br>[Proto]: $p<0.001$ |
| | | | | Paired two-sample bootstrap | [GFAP+]-[Proto]: 8.99 ± 2.07% | $p<0.0001$ |
| | | AUROC | 1000 random seeds | One-sample bootstrap hypothesis test against chance [illegible] | [GFAP+]: 75.71 ± 1.06%<br>[Proto]: 80.04 ± 0.98% | [GFAP+]: $p<0.001$<br>[Proto]: $p<0.001$ |
| | | | | Paired two-sample bootstrap | [GFAP+]-[Proto]: -4.33 ± 1.40% | $p=0.004$ |
| Figure 3d. Estimates are reported as mean ± SEM. | | | | | | |
| **Fig. 3** | **d 1st column** | Change in the degree of contextual encoding across days (1st block, Days 1-29) | 139 sessions from 5 mice | Repeated-measures correlation | r(133) = 0.1922 | $p=0.0255$ |
| | | Change in the degree of contextual encoding across the reversal (Days 29-30) | 10 sessions from 5 mice | One-sample Kolmogorov-Smirnov test to | D(5) = 0.2011 | $p=0.9599$ |
| | | | | Paired two-sample t-test | Day 29: 67.39 ± 12.81%<br>Day 30: 59.35 ± 4.50%<br>Days 29-30: -8.05 ± 16.00%,<br>t(4) = -0.5623 | $p = 0.6039$ |

| | | | | | | |
|---|---|---|---|---|---|---|
| | | Change in the degree of contextual encoding across days (2nd block, Days 30-46) | 82 sessions from 5 mice | Repeated-measures correlation | r(76) = 0.3980 | $p=3.07\times10^{-4}$ |
| | **d 2nd column** | Change in the degree of contextual encoding across days (1st block, Days 1-29) | 139 sessions from 5 mice | Repeated-measures correlation | r(133) = 0.6939, | $p=1.05\times10^{-20}$ |
| | | Change in the degree of contextual encoding across the reversal (Days 29-30) | 10 sessions from 5 mice | One-sample Kolmogorov-Smirnov test to | D(5) = 0.1895 | $p=0.9656$ |
| | | | | Paired two-sample t-test | Day 29: 88.35 ± 4.14%<br>Day 30: 36.76 ± 2.22%<br>Days 29-30: -51.59 ± 3.22%,<br>t(4) = -17.9183 | $p=5.7\times10^{-5}$ |
| | | Change in the degree of contextual encoding across days (2nd block, Days 30-46) | 82 sessions from 5 mice | Repeated-measures correlation | r(76) = 0.4093 | $p=1.98\times10^{-4}$ |
| | **d 3rd column** | Change in the degree of contextual encoding across days (1st block, Days 1-29) | 139 sessions from 5 mice | Repeated-measures correlation | r(133) = 0.1739 | $p=0.0437$ |
| | | Change in the degree of contextual encoding across the reversal (Days 29-30) | 10 sessions from 5 mice | One-sample Kolmogorov-Smirnov test to | D(5) = 0.1689 | $p=0.9735$ |
| | | | | Paired two-sample t-test | Day 29: 60.12 ± 17.32%<br>Day 30: 66.22 ± 10.47%<br>Days 29-30: 6.10 ± 27.28%,<br>t(4) = 0.2501 | $p=0.8148$ |

| | | | | | | |
|---|---|---|---|---|---|---|
| | | Change in the degree of contextual encoding across days (2nd block, Days 30-46) | 82 sessions from 5 mice | Repeated-measures correlation | r(76) = 0.2888 | *p*=0.0103 |
| | **d 4th column** | Change in the degree of contextual | 139 sessions from 5 | Repeated-measures correlation | r(133) = 0.6460 | $p=2.6\times10^{-17}$ |
| | | Change in the degree of contextual encoding across the reversal (Days 29-30) | 10 sessions from 5 mice | One-sample Kolmogorov-Smirnov test to | D(5) = 0.1328 | *p*=0.9874 |
| | | | | Paired two-sample t-test | Day 29: 86.09 ± 4.45%<br>Day 30: 40.85 ± 3.16%<br>Days 29-30: -45.25 ± 4.79%,<br>t(4) = -10.5721 | $p=4.52\times10^{-4}$ |
| | | Change in the degree of contextual encoding across days (2nd block, Days 30-46) | 82 sessions from 5 mice | Repeated-measures correlation | r(76) = 0.5597 | $p=9.98\times10^{-8}$ |
| | **e (ME, LNP, GFAP+)** | Correlation between the degree of contextual encoding (accuracy) and the behavioral optimality (factor) | 138 sessions from 5 mice | Repeated-measures correlation | r(132) = 0.1005 | *p*=0.2480 |
| | **e (ME, LNP, Proto)** | Correlation between the degree of contextual encoding (accuracy) and the behavioral optimality (factor) | 138 sessions from 5 mice | Repeated-measures correlation | r(132) = 0.6436 | $p=5.01\times10^{-17}$ |
| | **e (ME, LNP)** | Difference between correlation coefficients | 138 sessions from 5 mice | Paired two-sample permutation test | | *p*<0.001 |

| | | | | | | |
|---|---|---|---|---|---|---|
| | **e (ME, RNP, GFAP+)** | Correlation between the degree of contextual encoding (accuracy) and the behavioral optimality (factor) | 82 sessions from 5 mice | Repeated-measures correlation | r(76) = 0.1979 | *p*=0.0825 |
| | **e (ME, RNP, Proto)** | Correlation between the degree of contextual encoding (accuracy) and the behavioral optimality (factor) | 82 sessions from 5 mice | Repeated-measures correlation | r(76) = 0.3096 | *p*=0.0058 |
| | **e (ME, RNP)** | Difference between correlation | 82 sessions from 5 | Paired two-sample permutation | | *p*=0.3297 |
| | **f (MX, LNP, GFAP+)** | Correlation between the degree of contextual encoding (accuracy) and the behavioral optimality (factor) | 138 sessions from 5 mice | Repeated-measures correlation | r(132) = 0.0099 | *p*=0.9099 |
| | **f (MX, LNP, Proto)** | Correlation between the degree of contextual encoding (accuracy) and the behavioral optimality (factor) | 138 sessions from 5 mice | Repeated-measures correlation | r(132) = 0.6338 | $p=2.04\times10^{-16}$ |
| | **f (MX, LNP)** | Difference between correlation coefficients | 138 sessions from 5 mice | Paired two-sample permutation test | | $p< 0.001$ |

| | | | | | | |
|---|---|---|---|---|---|---|
| | **f (MX, RNP, GFAP+)** | Correlation between the degree of contextual encoding (accuracy) and the behavioral optimality (factor) | 82 sessions from 5 mice | Repeated-measures correlation | r(76) = 0.0982 | $p$=0.3923 |
| | **f (MX, RNP, Proto)** | Correlation between the degree of contextual encoding (accuracy) and the behavioral optimality (factor) | 82 sessions from 5 mice | Repeated-measures correlation | r(76) = 0.4271 | $p=9.61\times10^{-5}$ |
| | **f (MX, RNP)** | Difference between correlation coefficients | 82 sessions from 5 mice | Paired two-sample permutation test | | $p$=0.008 |
| **Fig. 4** | **c** | Two-way ANOVA | | Interaction | F (240, 1928) = 18.64 | *p<0.0001* |
| | | | | Raw Factor | F (240, 1928) = 17.23 | *p<0.0001* |
| | | | | Column Factor | F (1, 1928) = 213.2 | *p<0.0001* |
| | | Wilcoxon test | | VEH vs. C21 | W = -48 | *p=0.012* |
| | **e** | Two-way ANOVA | | Interaction | F (19, 160) = 174.6 | *p<0.0001* |
| | | | | Raw Factor | F (19, 160) = 23.83 | *p<0.0001* |
| | | | | Column Factor | F (1, 160) = 749.9 | *p<0.0001* |
| | **f** | Two-way ANOVA | | Interaction | F (19, 160) = 320.6 | *p<0.0001* |
| | | | | Raw Factor | F (19, 160) = 17.64 | *p<0.0001* |
| | | | | Column Factor | F (1, 160) = 1960 | *p<0.0001* |
| **Fig. 5** | **b** | Two-way ANOVA | | Interaction | F (12, 104) = 13.61 | *p<0.0001* |
| | | | | Raw Factor | F (12, 104) = 6.641 | *p<0.0001* |
| | | | | Column Factor | F (1, 104) = 1289 | *p<0.0001* |
| **Fig. 6** | **d** | Two-way ANOVA | | Interaction | F (12, 104) = 131.1 | *p<0.0001* |
| | | | | Raw Factor | F (12, 104) = 7.312 | *p<0.0001* |
| | | | | Column Factor | F (1, 104) = 487.3 | *p<0.0001* |
| | **e** | Two-way ANOVA | | Interaction | F (12, 104) = 221.8 | *p<0.0001* |
| | | | | Raw Factor | F (12, 104) = 58.98 | *p<0.0001* |
| | | | | Column Factor | F (1, 104) = 1007 | *p<0.0001* |
| **Fig. 7.** | **c** | Two-way ANOVA | | Interaction | F (12, 104) = 220.7 | *p<0.0001* |
| | | | | Raw Factor | F (12, 104) = 63.92 | *p<0.0001* |
| | | | | Column Factor | F (1, 104) = 1023 | *p<0.0001* |

| | | | | | |
|---|---|---|---|---|---|
| | **D** | Two-way ANOVA | Interaction | F (12, 104) = 113.7 | *p<0.0001* |
| | | | Raw Factor | F (12, 104) = 7.778 | *p<0.0001* |
| | | | Column Factor | F (1, 104) = 499.4 | *p<0.0001* |
| **ED Fig. 7** | **C21 #1** | Two-way ANOVA | Interaction | F (2, 24) = 11.89 | *p<0.0001* |
| | | | Raw Factor | F (2, 24) = 23.85 | *p<0.0001* |
| | | | Column Factor | F (1, 24) = 87.56 | *p<0.0001* |
| | **Saline** | Two-way ANOVA | Interaction | F (2, 24) = 4.086 | *p<0.0001* |
| | | | Raw Factor | F (2, 24) = 3.783 | *p<0.0001* |
| | | | Column Factor | F (1, 24) = 4.086 | *p<0.0001* |
| | **C21 #2** | Two-way ANOVA | Interaction | F (2, 24) = 28.10 | *p<0.0001* |
| | | | Raw Factor | F (2, 24) = 55.07 | *p<0.0001* |
| | | | Column Factor | F (1, 24) = 47.79 | *p<0.0001* |
| **ED Fig. 9** | **a** | One-way ANOVA | Specificity | F(2,12) = 0.5068 | *p=0.910* |
| | **b** | One-way ANOVA | Specificity | F(2,12) = 1.5230 | *p=0.779* |
| | **c** | One-way ANOVA | Specificity | F(2,12) = 0.5340 | *p=0.946* |
| | **d** | One-way ANOVA | Specificity | F(2,12) = 0.8103 | *p=0.666* |

**Supplementary Table 2. Statistical details for “Robustness of contextual encoding”**

| Figure | | Statistical Tests | | Comparison | Value & [illegible] | *P* value |
|---|---|---|---|---|---|---|
| Figure 3c. Estimates are reported as mean ± bootstrap standard error, as they are based on random under-sampling. [0.5 s time window] | | | | | | |
| **Fig. 3** | **c, ME** | Accuracy across blocks (Overall) Accuracy in the left context (LNP) | 1000 random seeds 1000 random seeds | One-sample bootstrap hypothesis test against chance level | [GFAP+]: 66.44 ± 1.17% [Proto]: 72.24 ± 1.10% | [GFAP+]: *p*<0.001 [Proto]: *p*<0.001 |
| | | | | One-sample bootstrap hypothesis test against chance level | [GFAP+]: 59.21 ± 2.46% [Proto]: 78.58 ± 1.21% | [GFAP+]: *p*<0.001 [Proto]: *p*<0.001 |
| | | Accuracy in the right context (RNP) AUROC | 1000 random seeds 1000 random seeds | One-sample bootstrap hypothesis test against chance level | [GFAP+]: 73.67 ± 2.15% [Proto]: 65.91 ± 1.76% | [GFAP+]: *p*<0.001 [Proto]: *p*<0.001 |
| | | | | One-sample bootstrap hypothesis test against chance level | [GFAP+]: 72.75 ± 1.14% [Proto]: 78.65 ± 1.01% | [GFAP+]: *p*<0.001 [Proto]: *p*<0.001 |
| | | Accuracy across blocks (Overall) Accuracy in the left context (LNP) | 1000 random seeds 1000 random seeds | One-sample bootstrap hypothesis test against chance level | [GFAP+]: 66.44 ± 1.17% [Proto]: 72.24 ± 1.10% | [GFAP+]: *p*<0.001 [Proto]: *p*<0.001 |
| | | | | One-sample bootstrap hypothesis test against chance level | [GFAP+]: 59.21 ± 2.46% [Proto]: 78.58 ± 1.21% | [GFAP+]: *p*<0.001 [Proto]: *p*<0.001 |
| | | Accuracy in the right context (RNP) AUROC | 1000 random seeds 1000 random seeds | One-sample bootstrap hypothesis test against chance level | [GFAP+]: 73.67 ± 2.15%, [Proto]: 65.91 ± 1.76% | [GFAP+]: *p*<0.001 [Proto]: *p*<0.001 |
| | | | | One-sample bootstrap hypothesis test against chance level | [GFAP+]: 72.75 ± 1.14% [Proto]: 78.65 ± 1.01% | [GFAP+]: *p*<0.001 [Proto]: *p*<0.001 |
| | **c, MX** | Accuracy across blocks (Overall) Accuracy in the left context (LNP) | 1000 random seeds 1000 random seeds | One-sample bootstrap hypothesis test against chance level | [GFAP+]: 61.97 ± 1.16%[Proto]: 62.31 ± 1.22% | [GFAP+]: *p*<0.001 [Proto]: *p*<0.001 |

| | | | | | | |
|---|---|---|---|---|---|---|
| | | | | One-sample bootstrap hypothesis test against chance level | [GFAP+]: 58.34 ± 1.87%<br>[Proto]: 65.75 ± 1.47% | [GFAP+]: *p*<0.001<br>[Proto]: *p*<0.001 |
| | | Accuracy in the right context (RNP)<br>AUROC | 1000 random seeds<br>1000 random seeds | One-sample bootstrap hypothesis test against chance level | [GFAP+]: 65.59 ± 1.82%<br>[Proto]: 58.88 ± 1.98% | [GFAP+]: *p*<0.001<br>[Proto]: *p*<0.001 |
| | | | | One-sample bootstrap hypothesis test against chance level | [GFAP+]: 65.89 ± 1.15%<br>[Proto]: 66.92 ± 1.21% | [GFAP+]: *p*<0.001<br>[Proto]: *p*<0.001 |
| | | Accuracy across blocks (Overall)<br>Accuracy in the left context (LNP) | 1000 random seeds<br>1000 random seeds | One-sample bootstrap hypothesis test against chance level | [GFAP+]: 61.97 ± 1.16%<br>[Proto]: 62.31 ± 1.22% | [GFAP+]: *p*<0.001<br>[Proto]: *p*<0.001 |
| | | | | One-sample bootstrap hypothesis test against chance level | [GFAP+]: 58.34 ± 1.87%<br>[Proto]: 65.75 ± 1.47% | [GFAP+]: *p*<0.001<br>[Proto]: *p*<0.001 |
| | | Accuracy in the right context (RNP)<br>AUROC | 1000 random seeds<br>1000 random seeds | One-sample bootstrap hypothesis test against chance level | [GFAP+]: 65.59 ± 1.82%<br>[Proto]: 58.88 ± 1.98% | [GFAP+]: *p*<0.001<br>[Proto]: *p*<0.001 |
| | | | | One-sample bootstrap hypothesis test against chance level | [GFAP+]: 65.89 ± 1.15%<br>[Proto]: 66.92 ± 1.21% | [GFAP+]: *p*<0.001<br>[Proto]: *p*<0.001 |
| Figure 3c. Estimates are reported as mean ± bootstrap standard error, as they are based on random under-sampling. [1 s time window] | | | | | | |
| **Fig. 3** | **c, ME** | Accuracy across blocks (Overall) | 1000 random seeds | One-sample bootstrap hypothesis test against chance level | [GFAP+]: 68.74 ± 1.18%<br>[Proto]: 75.26 ± 1.07% | [GFAP+]: *p*<0.001<br>[Proto]: *p*<0.001 |
| | | Accuracy in the left context (LNP) | 1000 random seeds | One-sample bootstrap hypothesis test against chance level | [GFAP+]: 60.87 ± 2.62%<br>[Proto]: 82.58 ± 1.13% | [GFAP+]: *p*<0.001<br>[Proto]: *p*<0.001 |
| | | Accuracy in the right context (RNP) | 1000 random seeds | One-sample bootstrap hypothesis test against chance level | [GFAP+]: 76.60 ± 2.21%<br>[Proto]: 67.95 ± 1.74% | [GFAP+]: *p*<0.001<br>[Proto]: *p*<0.001 |

| | | | | | | |
|---|---|---|---|---|---|---|
| | | AUROC | 1000 random seeds | One-sample bootstrap hypothesis test against chance level | [GFAP+]: 75.49 ± 1.08%<br>[Proto]: 82.84 ± 0.96% | [GFAP+]: *p*<0.001<br>[Proto]: *p*<0.001 |
| | **c, MX** | Accuracy across blocks (Overall) | 1000 random seeds | One-sample bootstrap hypothesis test against chance level | [GFAP+]: 66.28 ± 1.14%<br>[Proto]: 67.77 ± 1.16% | [GFAP+]: *p*<0.001<br>[Proto]: *p*<0.001 |
| | | Accuracy in the left context (LNP) | 1000 random seeds | One-sample bootstrap hypothesis test against chance level | [GFAP+]: 58.19 ± 1.82%<br>[Proto]: 73.89 ± 1.35% | [GFAP+]: *p*<0.001<br>[Proto]: *p*<0.001 |
| | | Accuracy in the right context (RNP) | 1000 random seeds | One-sample bootstrap hypothesis test against chance level | [GFAP+]: 74.38 ± 1.65%<br>[Proto]: 61.64 ± 1.76% | [GFAP+]: *p*<0.001<br>[Proto]: *p*<0.001 |
| | | AUROC | 1000 random seeds | One-sample bootstrap hypothesis test against chance level | [GFAP+]: 71.47 ± 1.10%<br>[Proto]: 72.81 ± 1.14% | [GFAP+]: *p*<0.001<br>[Proto]: *p*<0.001 |
| Figure 3c. Estimates are reported as mean ± bootstrap standard error, as they are based on random under-sampling. [1.5 s time window] | | | | | | |
| **Fig. 3** | **c, ME** | Accuracy across blocks (Overall) | 1000 random seeds | One-sample bootstrap hypothesis test against chance level | [GFAP+]: 69.81 ± 1.19%<br>[Proto]: 77.25 ± 1.04% | [GFAP+]: *p*<0.001<br>[Proto]: *p*<0.001 |
| | | Accuracy in the left context (LNP) | 1000 random seeds | One-sample bootstrap hypothesis test against chance level | [GFAP+]: 62.78 ± 2.52%<br>[Proto]: 84.35 ± 1.14% | [GFAP+]: *p*<0.001<br>[Proto]: *p*<0.001 |
| | | Accuracy in the right context (RNP) | 1000 random seeds | One-sample bootstrap hypothesis test against chance level | [GFAP+]: 76.83 ± 2.07%<br>[Proto]: 70.14 ± 1.70% | [GFAP+]: *p*<0.001<br>[Proto]: *p*<0.001 |
| | | AUROC | 1000 random seeds | One-sample bootstrap hypothesis test against chance level | [GFAP+]: 77.21 ± 1.03%<br>[Proto]: 84.81 ± 0.88% | [GFAP+]: *p*<0.001<br>[Proto]: *p*<0.001 |
| | **c, MX** | Accuracy across blocks (Overall) | 1000 random seeds | One-sample bootstrap hypothesis test against chance level | [GFAP+]: 68.35 ± 1.17%<br>[Proto]: 72.72 ± 1.06% | [GFAP+]: *p*<0.001 001<br>[Proto]: *p*<0.001 |

| | | | | | | |
|---|---|---|---|---|---|---|
| | | Accuracy in the left context (LNP) | 1000 random seeds | One-sample bootstrap hypothesis test against chance level | [GFAP+]: 60.05 ± 1.99%<br>[Proto]: 78.45 ± 1.30% | [GFAP+]: *p*<0.001<br>[Proto]: *p*<0.001 |
| | | Accuracy in the right context (RNP) | 1000 random seeds | One-sample bootstrap hypothesis test against chance level | [GFAP+]: 76.66 ± 1.56%<br>[Proto]: 66.98 ± 1.68% | [GFAP+]: *p*<0.001<br>[Proto]: *p*<0.001 |
| | | AUROC | 1000 random seeds | One-sample bootstrap hypothesis test against chance level | [GFAP+]: 73.75 ± 1.07%<br>[Proto]: 77.91 ± 1.03% | [GFAP+]: *p*<0.001<br>[Proto]: *p*<0.001 |
| Figure 3c. Estimates are reported as mean ± bootstrap standard error, as they are based on random under-sampling. [2 s time window] | | | | | | |
| **Fig. 3** | **c, ME** | Accuracy across blocks (Overall) | 1000 random seeds | One-sample bootstrap hypothesis test against chance level | [GFAP+]: 71.46 ± 1.12%<br>[Proto]: 77.80 ± 1.03% | [GFAP+]: *p*<0.001<br>[Proto]: *p*<0.001 |
| | | Accuracy in the left context (LNP) | 1000 random seeds | One-sample bootstrap hypothesis test against chance level | [GFAP+]: 64.95 ± 2.34%<br>[Proto]: 84.44 ± 1.26% | [GFAP+]: *p*<0.001<br>[Proto]: *p*<0.001 |
| | | Accuracy in the right context (RNP) | 1000 random seeds | One-sample bootstrap hypothesis test against chance level | [GFAP+]: 77.97 ± 1.83%<br>[Proto]: 71.16 ± 1.64% | [GFAP+]: *p*<0.001<br>[Proto]: *p*<0.001 |
| | | AUROC | 1000 random seeds | One-sample bootstrap hypothesis test against chance level | [GFAP+]: 78.88 ± 1.02%<br>[Proto]: 85.32 ± 0.83% | [GFAP+]: *p*<0.001<br>[Proto]: *p*<0.001 |
| | **c, MX** | Accuracy across blocks (Overall) | 1000 random seeds | One-sample bootstrap hypothesis test against chance level | [GFAP+]: 69.64 ± 1.13%<br>[Proto]: 74.89 ± 1.00% | [GFAP+]: *p*<0.001<br>[Proto]: *p*<0.001 |
| | | Accuracy in the left context (LNP) | 1000 random seeds | One-sample bootstrap hypothesis test against chance level | [GFAP+]: 61.25 ± 1.99%<br>[Proto]: 80.75 ± 1.22% | [GFAP+]: *p*<0.001<br>[Proto]: *p*<0.001 |
| | | Accuracy in the right context (RNP) | 1000 random seeds | One-sample bootstrap hypothesis test against chance level | [GFAP+]: 78.03 ± 1.54%<br>[Proto]: 69.04 ± 1.60% | [GFAP+]: *p*<0.001<br>[Proto]: *p*<0.001 |

| | | | | | | |
|---|---|---|---|---|---|---|
| | | AUROC | 1000 random seeds | One-sample bootstrap hypothesis test against chance level | [GFAP+]: 75.71 ± 1.06%<br>[Proto]: 80.04 ± 0.98% | [GFAP+]: $p<0.001$<br>[Proto]: $p<0.001$ |

**Supplementary Table 3. Statistical details for “Validation of behavioral optimality factor”**

| **Figure** | | **Statistical Tests** | | **Comparison** | **Value & [illegible]** | ***P* value** |
|---|---|---|---|---|---|---|
| Figure 3c. To assess the robustness of our behavioral optimality factor, we conducted a leave-one-out factor analysis. Each behavioral metric (session time, discrimination ratio, trajectory regularity) was iteratively excluded while factor analysis was performed on the remaining two. The correlation between the excluded metric and the extracted latent factor was then computed. To control for potential confounds related to experimental experience ("Day After Reversal"), we first removed its linear trend from each metric using a linear mixed model before applying the analysis. | | | | | | |
| **Fig. 3** | **c** | Hidden metric: session time | 260 sessions from 5 mice | Repeated-measures correlation | r(254) = 0.3766 | $p$=4.74×10$^{-10}$ |
| | | Hidden metric: discrimination ratio | 260 sessions from 5 mice | Repeated-measures correlation | r(254) = 0.5153 | $p$=9.06×10$^{-19}$ |
| | | Hidden metric: trajectory regularity | 260 sessions from 5 mice | Repeated-measures correlation | r(254) = 0.5051 | $p$=5.51×10$^{-18}$ |
